# Reliable *in silico* ranking of engineered therapeutic TCR binding affinities with MMPB/GBSA

**DOI:** 10.1101/2021.06.21.449221

**Authors:** Rory M. Crean, Christopher R. Pudney, David K. Cole, Marc W. van der Kamp

**Author notes:** Corresponding authors, (RMC), (MWvdK). Science for Life Laboratory, Department of Chemistry − Biomedicinska Centrum, Uppsala University, Box 576, S-751 23 Uppsala, Sweden (Rory M. Crean).

## Abstract

Accurate and efficient in silico ranking of protein-protein binding affinities is useful for protein design with applications in biological therapeutics. One popular approach to rank binding affinities is to apply the molecular mechanics Poisson Boltzmann/generalized Born surface area (MMPB/GBSA) method to molecular dynamics trajectories. Here, we identify protocols that enable the reliable evaluation of T-cell receptor (TCR) variants binding to their target, peptide-human leukocyte antigens (pHLAs). We suggest different protocols for variant sets with few (≤4) or many mutations, with entropy corrections important for the latter. We demonstrate how potential outliers could be identified in advance and that just 5-10 replicas of short (4 ns) MD simulations may be sufficient for reproducible and accurate ranking of TCR variants. The protocols developed here can be applied towards in silico screening during the optimization of therapeutic TCRs, potentially reducing both the cost and time taken for biologic development.

## Introduction

Computational methods to predict the binding affinities of protein-protein interactions (PPIs) that are sufficiently accurate, reliable and high throughput have clear potential for application towards the rational design of biologic drugs. Many approaches (all with many variations available) including free energy perturbation (FEP), umbrella sampling, molecular docking and machine learning have all been applied to predict or rank order PPI binding affinities.^1–4^ Here, we focused on the molecular mechanics Poisson Boltzmann/generalized Born surface area (MMPB/GBSA) approach,^5^ which combines conformational sampling using molecular dynamics (MD) simulations with empirical calculations on these snapshots to estimate the binding free energy. This approach can be thought of as sitting somewhere in between the more accurate but more computationally expensive FEP method, and less accurate but computationally cheaper methods like docking.^6^ This approach should only be relied on for relative binding affinities (*i.e.,* ΔΔ*G* not Δ*G*) to rank order a set of similar potential drug candidates.^6^ An advantage of the MMPB/GBSA approach is that it can be decomposed to obtain per-residue contributions to the binding energy, which we and others have used to identify key residues and interactions which drive protein-protein binding.^7–9^ The information obtained from this decomposition analysis can be used to inform (semi-)rational design efforts towards enhanced affinity and/or selectivity drug candidates.^8, 10^

MMPB/GBSA has been used and evaluated extensively for many applications, and it is clear that tuning of the parameters and protocols applied can give significant improvements in accuracy, with such tuning typically being system specific (see *e.g.* refs. ^6, 11–16^). With this in mind, we aimed to identify an MMPB/GBSA protocol that provides reliable and accurate relative binding free energies for a PPI of great interest in the field of immuno-oncology,^17^ T-cell receptor (TCR) peptide-human leukocyte antigen (pHLA) complexes (TCR-pHLA, **Figure 1**). The TCR-pHLA interaction is a vital component of the adaptive immune system, with the TCR ultimately responsible for selectively binding specific peptide sequences presented on the surface of cells by the HLA. For HLA Class 1 proteins (the focus of this study), the peptides in pHLA complexes are sourced from proteins digested inside the cell: each cell presents peptide fragments of its cellular proteins on the extracellular surface. In the natural immune system, TCRs can specifically identify antigenic peptide sequences on cells infected with pathogens, or expressing modified self-proteins in the case of cancer, presented on the cell surface by HLA molecules. TCR recognition of pHLA governs the activation of T cells that can lead to the direct killing and eradication of diseased cells.^18^

**Figure 1.**
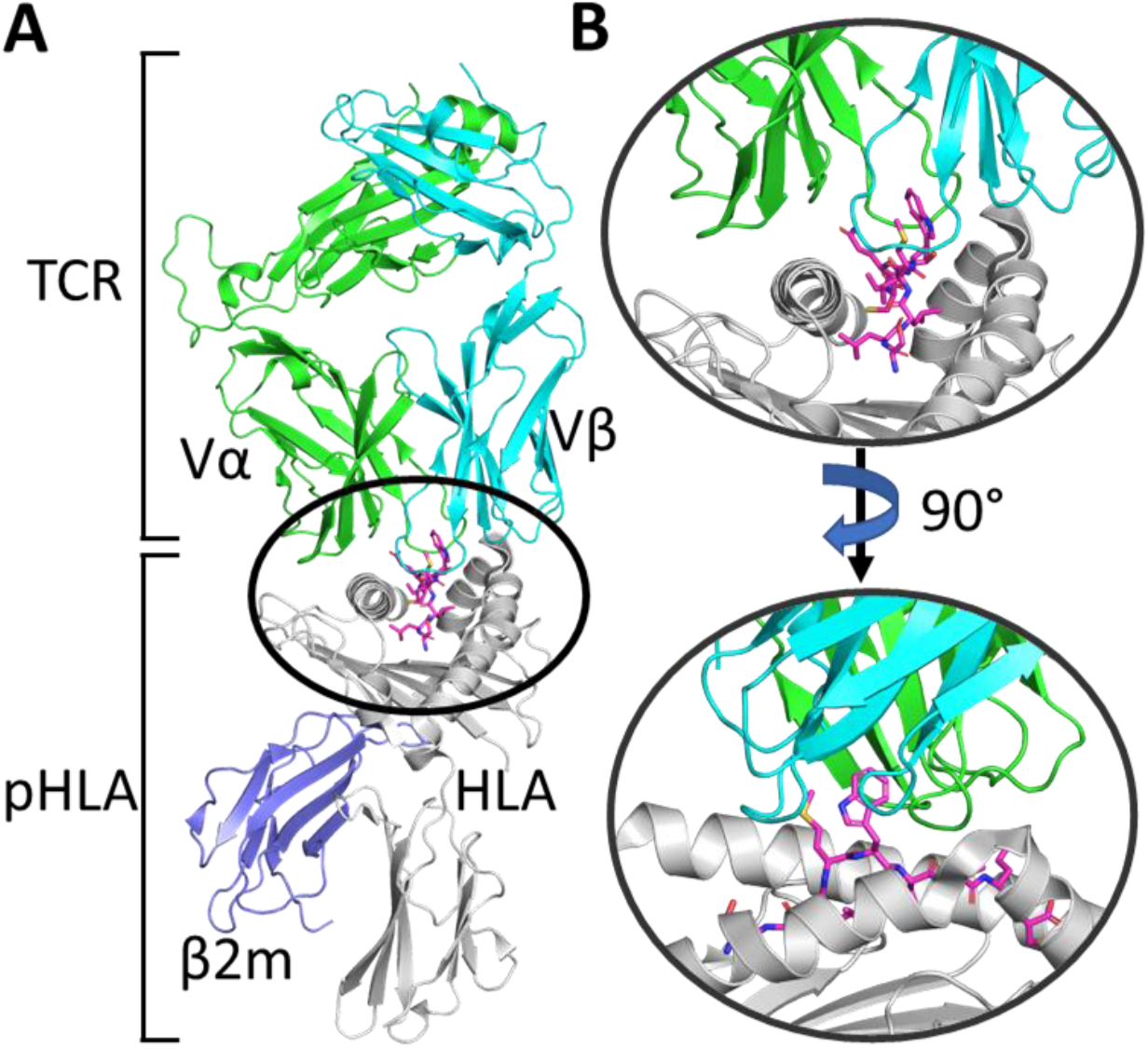
**A.** Overview of the TCR-pHLA complex. The T-cell receptor (TCR) is comprised of two (α and β) domains, which engage the peptide-human leukocyte antigen (pHLA) complex. **B.** Zoom in on the TCR-pHLA binding site from two different angles, demonstrating that the binding interface is composed of six complementarity-determining region (CDR) loops on the TCR, which engage both the peptide and two α-helices on the pHLA complex.

Affinity-enhanced, soluble, engineered TCRs are a class of therapeutic molecules which are designed to target a specific antigenic pHLA complex presented only by unhealthy (*e.g.* cancerous) cells, whilst simultaneously not binding the considerably large number of other pHLA complexes presented by “healthy” cells (in order to avoid off-target toxicity).^19^ This provides two deeply intertwined engineering challenges which must be addressed in order to produce a therapeutic TCR.^20^ That is, TCRs must have both strong affinity (natural TCRs have affinities in the ∼µM range,^21^ whilst therapeutic soluble TCRs are in the ∼pM range) and high specificity (to avoid the large number of off-targets). We have previously shown how both natural and engineered TCRs are able to achieve such specificity; through using a broad and energetically balanced network of interactions across the entire interface, making the TCR’s affinity very sensitive to mutations in either the peptide or HLA.^7^ Whilst most TCR affinity engineering studies reported in the literature have obtained affinity enhancement through experimental approaches (primarily those that utilize *in vitro* selection),^22–27^ docking^28, 29^ and structure based-rational design^30^ have also been successfully applied to engineer TCRs. Here, we envisage MMPB/GBSA as a technique that could be used to filter promising candidate mutations generated through a more high-throughput technique such as docking prior to experimental screening.

To date there has been no systematic study on how best to predict TCR-pHLA binding affinities using MMPB/GBSA, and herein, we aim to resolve this. To do this, we have evaluated a variety of MMPB/GBSA calculation protocols using two different TCR-pHLA test sets, one with 18 TCR variants with between 3-14 mutations (spread over most complementarity-determining region [CDR] loops) and one with 29 variants, of which 25 have just one mutation. The use of these two disparate test sets should allow us to identify a single protocol (if one exists) that works for both TCR-pHLA complexes and thus may be generally applicable for this biologically and therapeutically important protein-protein interaction.

## Methods

### Structure Preparation

X-ray crystal structures of the TCR-pHLA complexes of wild-type (WT) 1G4 and WT A6 were taken from PDBs 2BNR^31^ and 1AO7^32^ respectively, with the missing residues in PDB 1AO7 (located in the constant domain, away from the binding site) added in using PDB 4FTV,^33^ which has an identical (but resolved) constant domain to 1AO7. All simulations of point variants were performed using the WT structure, with mutations inserted using PyMOL^34^ (rotamers were selected based on recommendations from PyMOL v2.1, avoiding clashes as much as possible). Optimal His tautomerisation states and Asn and Gln side chain orientations were determined using MolProbity^35^, with all residues simulated in their standard protonation states at pH 7 (consistent with PROPKA 3.0^36^ predictions). His tautomerisation states were kept consistent between the WT and any variant structure simulated (see **Table S1** for tautomerisation states used). All structures were solvated in an octahedral water box such that no protein atom was within 10 Å of the box boundary, with the minimum number of either Na^+^ or Cl^−^ ions added as required to ensure total system neutrality. The 1G4 TCRs were solvated retaining the WT crystal structure waters, with any crystal water molecule that clashed with a newly inserted side-chain removed. For the A6 TCRs, the resolution of the WT structure (2.6 Å) is too low to identify (many) waters surrounding the binding site, so 3D-RISM^37, 38^ was used to calculate the radial distribution function (*g*(***r***)) of water surrounding the protein and the “Placevent” algorithm^39^ was used to solvate the protein based on the obtained *g*(***r***) (see **Supplementary Methods**), prior to solvation in a octahedral box.

### Molecular Dynamics Simulations

Molecular dynamics (MD) simulations were performed using GPU accelerated Amber16, ^40^ with the ff14SB^41^ force field and TIP3P water model were used to describe the protein and water molecules respectively. For each structure, a protocol of minimization, heating and equilibration (see **Supplementary Methods**) prior to production MD simulations in the NPT ensemble (298 K and 1 atm). For each structure, 25 replicas (with each replica assigned a different random velocity) of 4 ns long were performed, with the last 3 ns taken forward for MMPB/GBSA calculations. Simulations were performed with a 2 fs time step (with the SHAKE algorithm applied to all bonds containing hydrogen. The default 8 Å direct space non-bonded cut-off was applied with long range electrostatics evaluated using the particle mesh Ewald algorithm. Temperature and pressure regulation were performed using Langevin temperature control (collision frequency of 1 ps^−1^) and a Berendsen barostat (pressure relaxation time of 1 ps). Trajectory analysis was performed using CPPTRAJ.^42^ Hydrogen bonds (both solute-solute and water bridged) were considered formed if the donor-acceptor distances were less than 3 Å and the donor-hydrogen-acceptor angles were between 180±45°.

### MMPB/GBSA Theory and Methodology

The molecular mechanics generalized Born/Poisson–Boltzmann surface area (MMPB/GBSA) is an end state binding free energy calculation method which calculates the binding free energy (Δ𝐺_𝑏𝑖𝑛𝑑_) through the following equation:

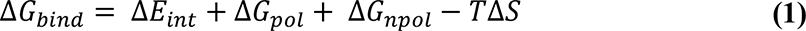

Where Δ𝐸_𝑖𝑛𝑡_ is the difference in the interaction energy, Δ𝐺_𝑝𝑜𝑙_ and Δ𝐺_𝑛𝑝𝑜𝑙_ are the polar and non-polar contributions to the solvation free energy respectively, and Δ𝑆 is the change in solute entropy. Δ𝐸_𝑖𝑛𝑡_ can be obtained directly from the force field energy terms:

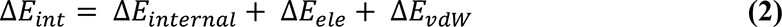

Where Δ𝐸_𝑖𝑛𝑡𝑒𝑟𝑛𝑎𝑙_ is the difference in the internal energy terms (i.e., bonding, angle, dihedral and improper torsions) and Δ𝐸_𝑒𝑙𝑒_ and Δ𝐸_𝑣𝑑𝑊_ are the electrostatic and Van der Waals (vdW) contributions. Note that in the single trajectory approach, which is used here, the contributions from Δ𝐸_𝑖𝑛𝑡𝑒𝑟𝑛𝑎𝑙_ cancel out. Δ𝐺_𝑝𝑜𝑙_ is obtained by solving either the Poisson Boltzmann (PB) or Generalized Born (GB) equations respectively. The non-polar contributions to the solvation free energy can be estimated from the solvent accessible surface area (SASA):

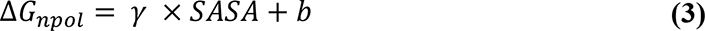

Where 𝛾 is surface tension (set to 0.00542 kcal mol^−1^ Å^−2^) and 𝑏 is an offset (set to 0.92 kcal mol^−1^).

Finally, 𝑇Δ𝑆 is an optional correction that accounts for the change in solute entropy. In this study we tested two different methods in order to calculate this, which are discussed in the section “Solute Entropy Corrections”.

For MMPB/GBSA calculations, frames were taken every 10 ps from the last 3 ns of each production MD simulation replica, meaning a total of 300 x 25 (number of replicas performed) frames were used for MMPB/GBSA calculations. Calculations were performed using the MPI version of MMPBSA.py^5^, with the GB-Neck2^43^ (*i.e.,* igb=8) solvent model for GBSA calculations, and the default PB solvent model for MMPBSA calculations. MMPB/GBSA calculations were performed with an implicit salt concertation of 150 mM (to match experimental assay conditions). In PBSA, the interior dielectric of the solute was varied (using the “indi” flag) as indicated in the text. (This is not an option for GBSA, where there is no interior dielectric value, as this is approximated through the use of Born radii.)

### Solute Entropy Corrections

The MMPB/GBSA approach does not account for the rigidification of the solutes upon binding (*i.e.,* the change in solute entropy contribution upon binding). We applied two different methods to predict a “correction” for this effect to the calculated binding free energies, using both the interaction entropy (Int-Entropy^44^) and the truncated normal mode analysis^45^ (Trunc-NMA) methods.

The Int-Entropy approach developed by Duan et al.^44^ uses the fluctuation of the gas phase contributions to Δ𝐺_𝑏𝑖𝑛𝑑_ (referred to as the interaction energy, Δ𝐸_𝑖𝑛𝑡_) to provide an estimate of 𝑇Δ𝑆. **Equation 2** shows how to calculate Δ𝐸_𝑖𝑛𝑡_. The per frame fluctuation of Δ𝐸_𝑖𝑛𝑡_ can then be determined by:

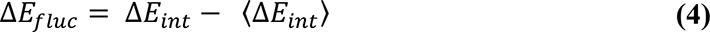

Where 〈Δ𝐸_𝑖𝑛𝑡_〉 is the ensemble average of Δ𝐸_𝑖𝑛𝑡_. Finally, 𝑇Δ𝑆 can be determined by:

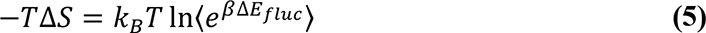

Where 𝛽 is 1⁄𝑘_𝐵_𝑇. For each different MMPBSA or MMGBSA calculation, we took the Δ𝐸_𝑖𝑛𝑡_ values obtained from all 7500 frames per complex and used this to calculate −𝑇Δ𝑆.

Normal mode analysis (NMA) uses vibrational frequency calculations of energy minimized structures of each state to determine the change in solute rigidity upon ligand binding and can therefore be used to determine 𝑇Δ𝑆 in **Equation 1**. To reduce the computational cost and noise associated with this approach, we used a modified version of this approach referred to as truncated-NMA (Trunc-NMA), developed by Kongsted and Ryde.^45^ In Trunc-NMA, only a subset of atoms located near the binding site are used for the entropy calculation. Residues located close to the binding site are treated as flexible (*i.e.,* allowed to move and therefore contribute to a vibrational frequency calculation), whilst residues further away from the binding site are included in a “buffer zone” and held fixed throughout the minimization and vibrational frequency calculations. Further, water molecules that surround the binding site are also often included as part of the buffer region. In the Trunc-NMA approach used here (see **Supplementary Methods** for further details), we retained all receptor (pHLA) residues within ∼16 Å of any ligand (TCR) residue and vice versa, using the WT crystal structure to determine distances. Any breakages introduced into the sequence were acetylated or amidated, using the coordinates from the first deleted residue. Those residues within the range 12-16 Å were kept frozen for both the optimization and vibrational frequency calculations. A shell of 1000 water molecules were also retained (and kept frozen throughout) around the binding site. For the frequency calculations of the free ligand or receptor, 500 water molecules were included for each structure. Energy minimization was performed using sander (Ambertools18^40^) with GB implicit solvent and performed until the RMSD was less than 10^−6^ kcal mol^−1^ Å^−1^. Frequency calculations were performed *in vacuo* using a modified version of the Nmode program (from Amber14), to allow use of the “ibelly” command, which allows for the freezing of atoms during the energy minimization and vibrational frequency calculations. Frozen atoms therefore have no (direct) impact on the entropy estimates obtained. The Trunc-NMA approach was only applied to the 1G4 set of TCRs and was performed on frames taken every 100 ps from the last 3 ns of each of the 25 replicas (750 frames per complex).

### Assessment of the Quality of Prediction

Experimentally determined ΔΔ𝐺’s (obtained from prior studies^26, 29, 33, 46^, see **Tables S2** and **S3** for affinities of all TCR-pHLA complexes studied) were compared to the computationally derived ΔΔ𝐺’s and assessed using the Pearson’s r (rp) value, Spearman’s rank (rs) and mean absolute deviation (MAD). These metrics were chosen as rp determines how linearly correlated the two data sets are, whilst rs assesses how monotonic the two data sets are (*i.e.,* how well do the computational results correctly rank order the experimental results). The MAD determines the average size of each residual from the linear fit. Error values associated with individual ΔΔ𝐺_𝑏𝑖𝑛𝑑_ calculations are the standard deviation obtained from the 25 replicas performed per complex. Bootstrapping with random replacement was performed using the R software package. In all instances, 1 million bootstrap resamples were constructed from the original 25 replicas performed per complex. Each resample was then used to calculate Spearman’s rank and Pearson correlation coefficient *r*, with the average values and 95% confidence intervals determined for different numbers of replicas.

### Simulation Timings

A single, 4 ns long MD simulation of a NVIDIA Pascal P100 GPU takes approximately 6 hours for a TCR-pHLA complex solvated in a water box (∼150,000 atoms). MMGB/PBSA calculations were performed on one dual socket Intel Ivybridge nodes with E5-2650v2 processors (clock rate 2.6GHz, 8 cores). To run MMGBSA and MMPBSA calculations on 300 frames (effectively one simulation run) took approximately 7 minutes and 60 minutes respectively. The above timings were not significantly affected by the addition of explicit waters. Trunc-NMA calculations were performed on one Intel SandyBridge node (16 cores with a 2.6 GHz clock rate). Int-Entropy calculations on 30 frames (effectively one simulation run) took approximately 4 hours.

## Results and discussion

In this study, we assessed the capability of MMPB/GBSA calculations to reproduce TCR-pHLA binding affinity relationships on two different test sets. The first (1G4) test set, was composed of 18 TCR variants all containing between 3-14 mutations from the WT, with mutations spread between 1-5 CDR loops.^26, 46^ In contrast, the second (A6) test set was composed of 29 TCR variants, with 25 of these being single point variants and the remaining 4 baring between 2-4 mutations.^29, 33^ The names 1G4 and A6, assigned to the WT TCR-pHLA complexes in their original publications (ref. 31 for 1G4 and ref. 32 for A6, respectively) will be used throughout the manuscript to define each system. We note that previously, we have used simulation and MMPBSA analysis (including decomposition of binding energies) to investigate, amongst others, four high affinity (affinity-enhanced) 1G4 variants and one A6 variant compared to their wild-type TCRs.^9^ This revealed that there are typically many TCR-peptide contacts and the peptide can contribute significantly to the overall TCR-pHLA binding affinity. Changes to the TCR-peptide interactions and its contributions, however, are typically modest, with the mechanisms of affinity enhancement being complex, often resulting from indirect and compensatory effects. The aim here was to identify protocols for affinity prediction (based on WT X-ray structures) that are not only reliable and reproducible, but also work well for the two disparate TCR test sets studied. With this in mind, we built on recommendations of others^15, 47–50^ in our use of many replicas of short MD simulations to obtain snapshots for MMPB/GBSA calculations. This “ensemble” based approach has been shown to outperform single or few replica simulations of much longer length, both in terms of reliability and predictability.^47, 49^ Specifically, we performed 25 independent MD simulations of 4 ns long, and used frames collected every 10 ps from the last 3 ns of each as input for MMPB/GBSA calculations (meaning a total of 7500 frames were used per TCR-pHLA complex). The prediction accuracy was assessed using the Pearson’s r (rp), Spearman’s rank (rs) and mean absolute deviation (MAD). These metrics were chosen as the rp measures the linear correlation between experiment, the MAD measures the average residual from the linear fit and the rs asses the ability to rank order binding affinities (arguably the rs is the most important metric in a design context).

### Modulation of the internal dielectric constant drastically improves predictability

For both the 1G4 and A6 TCRs test sets investigated, we assessed the ability of both MMPBSA and MMGBSA to predict relative binding affinities (**Figure 2**). Further, given previously reported successes at improving the quality of prediction for other systems,^6, 51–54^ we assessed the benefit of modifying the internal dielectric constant (𝜖_𝑖𝑛𝑡_) for the MMPBSA calculations.

**Figure 2.**
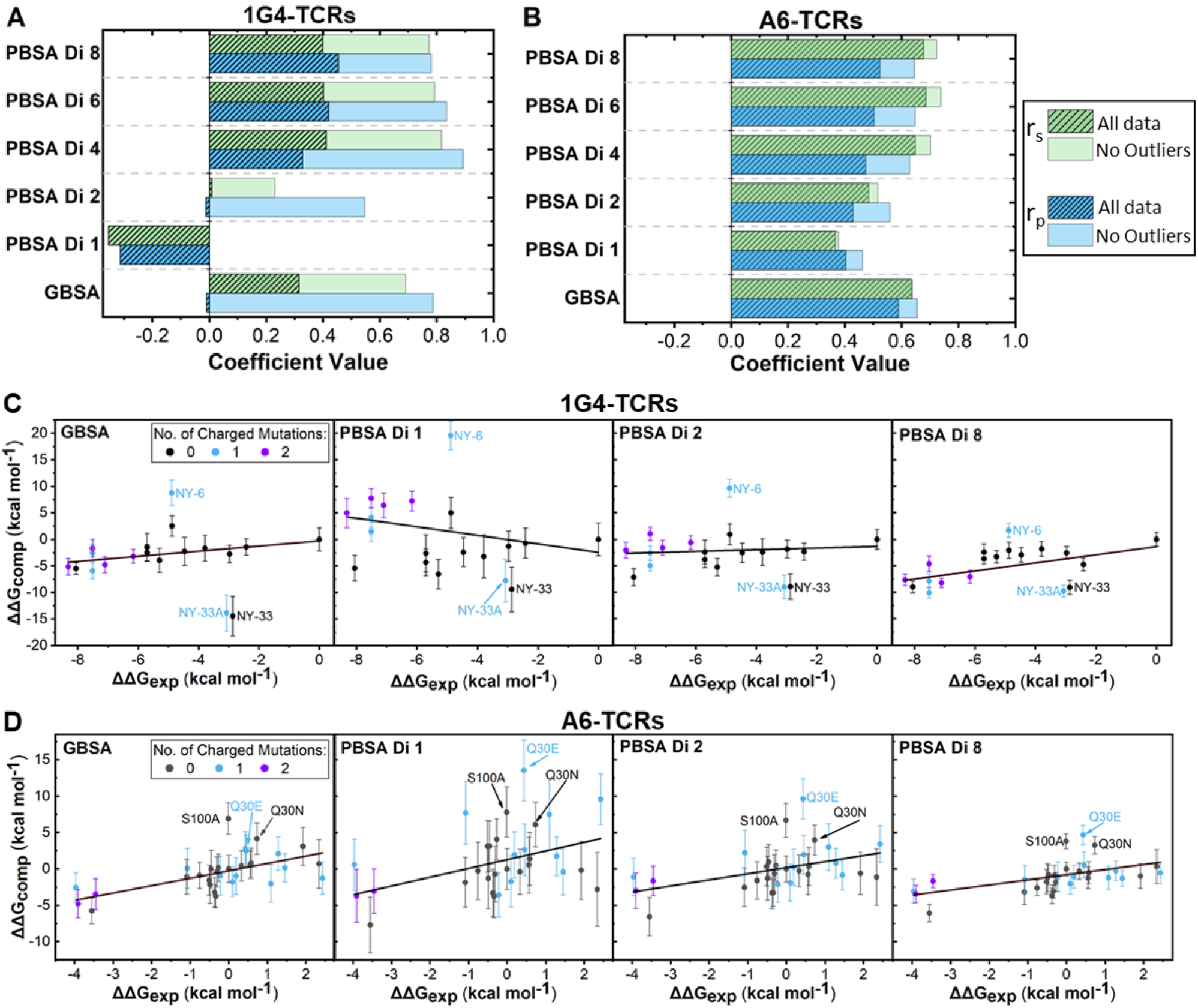
Modulation of the interior dielectric constant improves MMPBSA predictability. **A+B.** Determined Spearman’s rank (rs) and Pearson’s r (rp) values for MMPB/GBSA calculations for the 1G4 (**A**) and A6 (**B**) test sets. Results are plotted with and without the three identified outliers described in the text for both data sets. **C+D.** Exemplar scatter plots with lines of best fit for the 1G4 (**C**) and A6 (**D**) test sets using either MMGBSA or MMPBSA (at different internal dielectric constants) methodology. For **C**+**D**, outliers are labelled. Scatter plots in panels **C**+**D** are also colored according to the number of charged mutations made between the variant and the WT. Complete scatter graphs for all results are provided in Figures S1 and S2.

First, in the 1G4 test set, and to a lesser extent the A6 test set, increasing the 𝜖_𝑖𝑛𝑡_ used in the MMPBSA calculations progressively improved the prediction quality, with the effect largely flattening out for internal dielectric constants in the range of 4-8 (**Figure 2A+B** and **Table S5**). We also note that the standard deviations (SD) obtained from the 25 replicas for individual ΔΔ*G* measurements reduce as 𝜖_𝑖𝑛𝑡_ increases, with this effect again flattening out for 𝜖_𝑖𝑛𝑡_ values between 4-8 (**Figure 2C+D** and **Figures S1** and **S2**). For example, the average SD reduces from 2.8 to 1.3 kcal mol^−1^ and 3.5 to 1.7 kcal mol^−1^ when 𝜖_𝑖𝑛𝑡_ was increased from 1 to 4 for the 1G4 and A6 TCR systems, respectively. This data suggests that fewer replicas per variant may be required to obtain converged results when a higher 𝜖_𝑖𝑛𝑡_ value is used. Interestingly, for both test sets, the GB solvent model significantly outperformed the PB solvent model (at an 𝜖_𝑖𝑛𝑡_ of 1). This is perhaps surprising given that the GB solvent model is designed to reproduce the PB solvent model with an 𝜖_𝑖𝑛𝑡_ of 1.^43^ Although the majority of computational resource was spent on running the explicit solvent MD simulations for generating the conformational ensembles, it is worth noting that the MMGBSA method is approximately 8 times faster than MMPBSA (see **Methods** section “*Simulation Timings*” for further details). Its poor performance on the 1G4 test set, however, indicates that MMGBSA cannot be relied on for all TCR-pHLA combinations and should thus be compared to MMPBSA in the first instance.

It is challenging to provide a concrete answer as to the reason why increasing the 𝜖_𝑖𝑛𝑡_ can improve the quality of prediction, and why the 1G4 test set is more sensitive to this effect than the A6 test set. A recent MMPBSA study focused on predicting the correct binding pose for PPIs observed a weak relationship between the polar buried area (PBA) and the optimal 𝜖_𝑖𝑛𝑡_ to use.^51^ Systems with increasing PBA were recommended higher 𝜖_𝑖𝑛𝑡_ values, and based on the PBA of the WT TCR-pHLA complexes studied here (1310 Å^2^ and 1250 Å^2^ for WT-1G4 and WT-A6 respectively, determined using the COCOMAPS webserver^55^) a 𝜖_𝑖𝑛𝑡_ of approximately 2-4 would be recommended. Further, several MMPBSA alanine scanning studies have found the use of 𝜖_𝑖𝑛𝑡_ values greater than 1 to greatly improve the quality of prediction for the exchange of charged residues.^16, 56–59^ Finally, a recent study using a modified form of MMPBSA showed substantial improvement towards predicting the binding affinity for protein-protein interactions compared to the traditional MMPBSA approach.^60^ This modified form of MMPBSA considers the screening effect of ions on the electrostatic interactions between atoms and was found to be particularly beneficial in the case of highly charged systems. To assess the possibility that the outliers observed in the MMPBSA calculations with an 𝜖_𝑖𝑛𝑡_ of 1 were induced by changes in the charge of the TCR, we colored variants in **Figure 2C+D** according to the total number of charged mutations made from the WT. The benefit that increasing 𝜖_𝑖𝑛𝑡_ has on charged variants is clear for both data sets, but particularly striking for the 1G4 test set, as several affinity-enhanced variants (with ΔΔ*G* values < −6 kcal mol^−1^) are progressively reordered from some of the lowest affinity variants to some of the highest affinity variants.

For the 1G4 TCRs, three apparent outliers can be identified even at higher 𝜖_𝑖𝑛𝑡_ values or when using the GBSA approach (**Figure 2C**), and their negative impact is clear when comparing the prediction quality with and without the outliers included (**Figure 2A**). Their designation as outliers was validated by analysis of the residuals from linear regression between calculated and experimental binding affinity differences (**Figure S3**). Analysis of the CDR loop sequences of these TCR variants (**Figure 3A**) shows 5 mutations are made in their CDR3α loop which are not present in any of the other variants studied here (see **Table S4** for all sequences used). These differences in the CDR3α loop could therefore explain why these variants are outliers in the above data set. That is, these mutations may have notably altered the conformational dynamics/sampling of the loop (and/or neighboring regions), and this would likely not be accounted for by the short MD simulations (which start using the same backbone crystal structure as described in the **Methods**) performed here. This may be especially true in the case of NY-6, as its CDR3α loop contains both a mutation to remove a proline and another mutation to add a proline. In cases such as these, approaches that attempt to sample for changes in the TCR loop conformations upon mutation (such as those in refs. 61 or 62) could be used to generate the starting structures for MD simulations. Alternatively, there could be a significant change in the rigidity of the CDR3α-loop, such that the contribution from changes in solute entropy upon binding cannot be ignored for accurate ranking of these variants.

**Figure 3.**
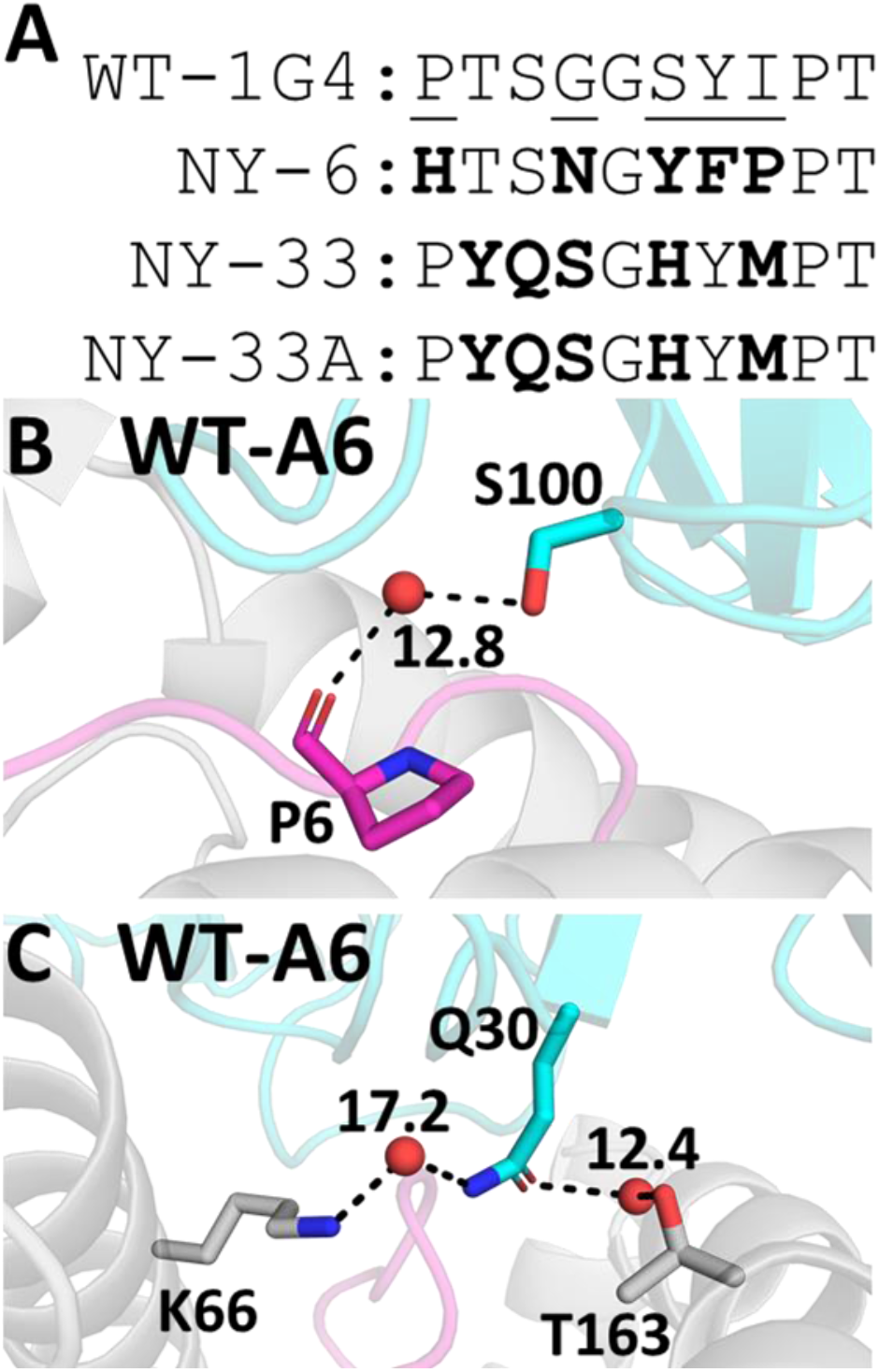
A potential rational for outliers identified in our MMPB/GBSA Calculations. **A.** Sequences of the CDR3α loop of the three 1G4 outliers, with positions mutated shown in bold. All 1G4 variant sequences are provided in **Table S4 B+C.** WT A6 TCR-pHLA structure with the two outlier mutation sites S100 (**B**) and Q30 (**C**) labelled. Predicted water sites (using 3D-RISM37,38 and Placevent39, see **Methods**) which form bridged water hydrogen bonds to pHLA residues are shown (here, all donor-acceptor heavy atom distances are within 3 Å). The calculated water density distribution function *g*(***r***) is shown for the water molecules, demonstrating that they are all predicted to have a very high occupancy.

For the A6 TCRs, 3 single point variants (Q30E, Q30N and S100A on the TCR α-chain) were consistently underestimated (**Figure 3D**). As we did for the 1G4 TCR test set above, residuals from linear regression were calculated, which supports our designation of these 3 single point variants as outliers (**Figure S4**). Our 3D-RISM calculations on WT A6 (used to solvate the protein due to the lack of water molecules available in the X-ray structure, see **Methods**) predicted strong affinity bridging water molecule sites at both of the above mutation sites in the WT A6-TCR (**Figure 3B+C**), with the 3D-RISM distribution function (*g*(***r***)) for water oxygen atoms calculated to be > 10 (note that the *g*(***r***) of bulk water is by definition 1). Further, both mutated side chains are predicted to make water-bridged hydrogen bonds with pHLA residues (**Figure 3B+C**). Specifically, HLA residues K66 and T163 for Q30 and peptide residue P6 for S100). Taken together, our data suggests that the outlier mutations may be poorly described due to not explicitly describing key solvent meditated hydrogen bonds through the use of an implicit solvent model (PB or GB) in our calculations.

Given the above observations, in the following sections we aimed to try to correct the outliers observed in both data sets and improve our overall prediction accuracy. We did this by (1) including explicit water molecules into our MMPB/GBSA calculations, and (2) introducing a correction for the change in solute entropy. Further, we note that our primary aim was to identify an approach that is ideally suitable for all TCR-pHLA complexes. It was therefore important to assess whether the inclusion of explicit water molecules and entropic corrections could have a deleterious effect on the overall quality of prediction (*i.e.,* through the introduction of an additional source of error and/or noise).

### Effect of Inclusion of Explicit Water Molecules

The inclusion of explicit water molecules has shown mixed success in the context of MMPB/GBSA calculations.^11, 14, 63–65^ When including explicit water molecules for calculating protein-small molecule binding affinities, common practice is to include the “X” closest water molecules to the ligand and retain these water molecules for the receptor calculation (as well as the complex calculation). In contrast, for a PPI, there are many possible ways to define which water molecules should be kept in the calculation and further, whether these waters are retained on the receptor or the ligand or some combination of both. Here, we took the X (where X is 10, 20, 30 or 50) closest waters to any oxygen or nitrogen atom on a selection of binding site residues located on the pHLA, and included these waters as part of the pHLA (*i.e.,* receptor) calculation, as well as the complex calculation (see **Supplementary Methods** for further details). We choose to keep all waters on the pHLA over a combination of the TCR and pHLA to ensure that all retained waters were close to a protein atom in both the bound and unbound MMPB/GBSA calculations. Given the results obtained for different solvent models and dielectric constants as described in **Figure 2**, we assessed the benefit of including explicit water molecules using both the MMGBSA and MMPBSA methods, setting 𝜖_𝑖𝑛𝑡_ to 6 for the MMPBSA calculations (**Figure 4**).

**Figure 4.**
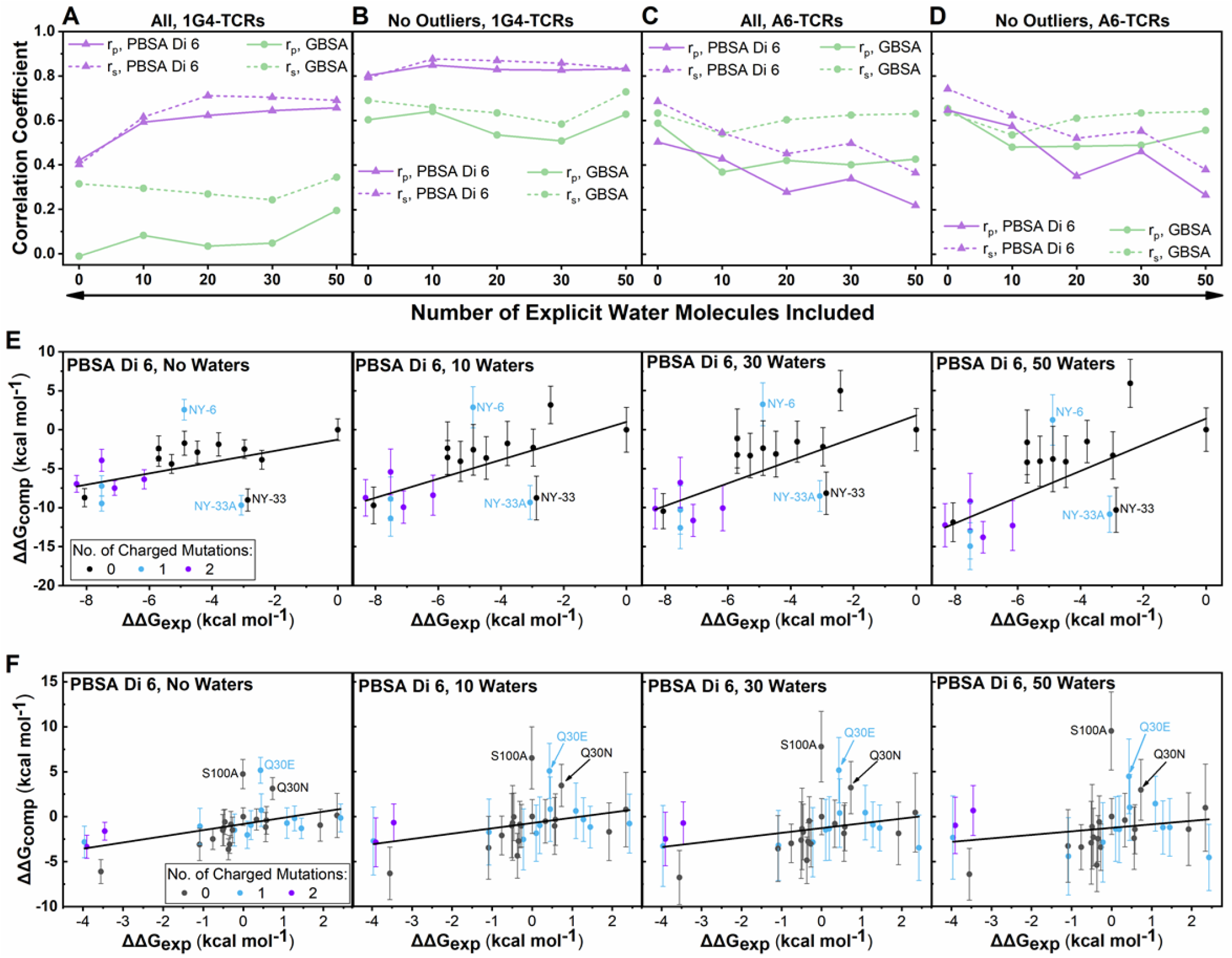
Impact of explicit water molecules on binding affinity predictions. **A-D**. Determined Spearman’s rank (rs) and Pearson’s R (rp) values for MMPB/GBSA calculations on the 1G4 (**A**+**B**) and A6 (**C**+**D**) test sets for different numbers of explicit water molecules included in the calculation. **E+F**. Exemplar scatter plots for the 1G4 (**E**) and A6 (**F**) test sets showing the impact of the inclusion of an increasing number of explicit water molecules when using the MMPBSA method with 𝜖𝑖𝑛𝑡 set to 6. Scatter points are colored according to the number of charged mutations made between the variant and the WT. Complete scatter graphs for all results are provided in **Figures S5-S8**.

Focusing first on the 1G4 set of TCRs, a beneficial effect was observed when explicit water molecules were included in the MMPBSA calculation with 𝜖_𝑖𝑛𝑡_ set to 6 for the entire dataset (**Figure 4A**). The prediction quality is only marginally improved when the outliers were excluded (**Figure 4B**), suggesting the inclusion of explicit water molecules helped improved these outlier data points. This additional benefit appears to be largely due to correctly ranking the highest affinity TCR variants (those with ΔΔ*G*exp < −6 kcal mol^−1^). Further, most of the beneficial effect of including explicit water molecules was observed after only 10 waters are included, with the improved prediction quality remaining fairly stable with increasing numbers of waters included. This observation of a lack of sensitivity to differing numbers of explicit waters is reassuring to note (as it is impossible to know the optimal number of waters to include *a priori*). However, adding explicit water molecules to the A6 test set negatively impacted the prediction accuracy, especially for the MMPBSA calculations (**Figure 4C**+**D, Table 6**). Notably, no X-ray crystal waters were used for this test set, which may in part explain the poor performance.

In contrast to the MMPBSA calculations, the inclusion of explicit water did not significantly improve the correlations for the MMGBSA approach. It should be noted, however, that the inclusion of explicit solvent increased the standard deviation obtained for the individual affinity estimates (**Figure 4E+F** and **Figures S5** and **S6**). For both the MMPBSA and MMGBSA simulations of the 1G4 test set, this increased deviation is partially compensated for by sampling a larger range of affinities. For instance, the MMPBSA ΔΔ*G*calc values vary by up to 12.2 kcal mol^−1^ for calculations with no water as compared to up to 20.9 kcal mol^−1^ for calculations with 50 waters included (with the three outliers described above removed, the variations for no water or 50 waters are 9.5 and 20.9 kcal mol^−1^, respectively). This was also reflected in the mean absolute deviation (MAD) values obtained (**Table S6**), in which increasing the number of water molecules consistently increased the MAD for both the 1G4 and A6 test sets. Although this observed increase in the MAD would be of concern if the ultimate goal is prediction of absolute binding affinity differences, it does not directly affect the rank ordering of candidate mutations (e.g., for design).

In contrast, the range of ΔΔ*G*calc values obtained for the A6 TCRs did not change significantly with increasing numbers of waters (**Figure 4F**, **Figures S9** and **S10**), indicating that the impact of the increased standard deviations observed may be particularly detrimental to the prediction accuracy for the A6 test set (as this therefore implies increased noise in the dataset).

As the A6 TCR data set consists primarily of single point mutations, whist the 1G4 set is composed entirely of multi-point variants, it is important to consider how significant the contribution of explicit water molecules is in describing the differences in affinity between variants (*i.e.,* ΔΔ*G* not Δ*G*). That is, mutations that do not (significantly) interrupt the solvation environment between the TCR and pHLA may not require explicit solvation in order to correctly rank their relative affinities, and instead the increased noise associated with the calculation may just worsen the prediction quality. One would expect single point mutations to significantly disrupt the water network less often than the multi-point mutants present in the 1G4 test set, which is consistent with our observations shown in **Figure 4**. Further, the A6 TCR model was solvated based on 3D-RISM calculations, as no crystallographic waters were resolved (due to the low resolution of the structure). This may therefore also provide a source of error, if any key binding site water molecules were incorrectly placed.

To try to identify how the binding site solvation environment may have changed for TCR-pHLA complexes with different TCR variants, we calculated the total average number of bridged water hydrogen bonds (Hbonds) as well as solute-solute Hbonds formed between the TCR and pHLA during our MD simulations (**Figure S9**). Whilst in the A6 data set we did observe a notable decrease in the average number of bridged water Hbonds for the Q30E variant (one of the outliers described above) as compared to the WT, other variants showed largely similar values, consistent with a largely unchanged binding site water network. We also note that the 1G4 outliers NY-33 and NY-33A had the largest number of solute-solute Hbonds (**Figure S9**), approximately 3 more Hbonds than most of the rest of the 1G4 test set. Our binding affinity calculations overestimated these outliers’ affinities (**Figures 2** and **4**, where no solute entropy correction term has yet been considered). This could suggest that enthalpy-entropy compensation is important for correctly ranking these outliers.^66^ That is, with additional Hbonds between the TCR and pHLA, one may expect a more favorable binding enthalpy term, which could be offset to a large degree by a less favorable binding entropy term. There are not enough data points, however, to determine if this is a general trend; we further note that outlier NY-6 did not show an increase in solute-solute hydrogen bonds, indicating outliers may also be caused through other effects.

### Impact of Solute Entropy Corrections

We evaluated two different methods to determine the change in solute entropy (𝑇Δ𝑆). The first is a modified version of the normal mode analysis (NMA) approach. In this approach, snapshots from MD are subjected to energy minimization and vibrational frequency calculations to obtain an estimate of the configurational entropy for each state. This approach is often not used in MMPB/GBSA applications due to its sizable computational cost and the large standard deviations obtained, which can often worsen the prediction quality.^6, 67^ However, Kongsted and Ryde introduced a modified approach whereby NMA is performed on a truncated region around the binding site, with a “buffer” region of amino acids and water molecules fixed in place to stabilize the conformation of the structure (**Figure 5**).^45^ This approach, referred to as truncated-NMA (Trunc-NMA) has been demonstrated to significantly reduce the computing time associated with the calculations, as well as reducing the magnitude of the error values obtained.^45, 68, 69^ Given that we did not expect entropy corrections to improve predictions for the A6 test set with (primarily) single point mutations, alongside the substantial computational cost of Trunc-NMA, we applied this approach on only the 1G4 set of TCRs. Even with this truncated approach, the time taken to run Trunc-NMA calculations was substantially greater than that for the standard MMPB/GBSA calculations (see the **Methods** section “Simulation Timings” for further details). This approach is thus not suitable for efficient, high-throughput screening of large numbers of variants.

**Figure 5.**
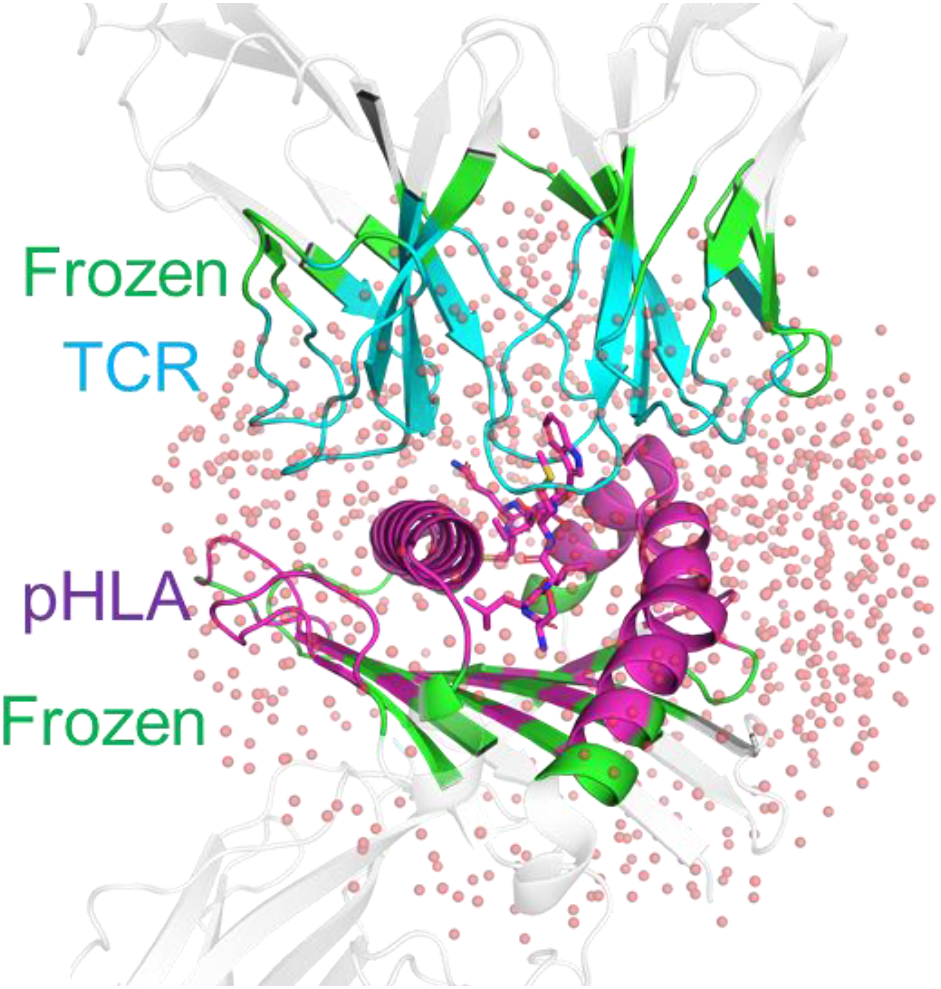
Illustration of the truncated-normal mode analysis (Trunc-NMA) method used to calculate a solute entropy correction for the 1G4 test set. Residues included in the Trunc-NMA calculations are colored in blue (TCR) or magenta (pHLA) if they are flexible in the NMA calculations or green if they are frozen (and therefore make up part of the buffer region). Residues colored in white are not included in the calculation (see **Methods**). The 1000 water molecules retained in the calculation are shown as transparent spheres.

The second method evaluated is known as the interaction entropy^44^ (Int-Entropy) approach, which determines the solute contributions to −𝑇Δ𝑆 from the fluctuations in the change in the gas phase interaction energy (*i.e.,* larger average fluctuations result in a larger value of −𝑇Δ𝑆, see **Methods** for further details). This approach has the advantage of not requiring additional simulations (as fluctuations of the gas phase interaction energy can be taken directly from the original MMPB/GBSA calculations) and has shown great promise as a correction for protein-ligand binding free energy calculations.^44, 54, 70, 71^

For both test sets, there was a clear reduction in the quality of prediction when the Int-Entropy corrections are applied (**Figure 6**). Analysis of individual scatter plots with and without this approach included (**Figure 6B+D**) illustrate that the Int-Entropy approach had a negative effect on the prediction accuracy. We note that the error bars plotted for calculations with the Int-Entropy approach do not include an estimate of the uncertainty of the Int-Entropy correction itself (as all frames are combined for a single estimate). Nevertheless, it is clear that the noise and/or error induced from the Int-Entropy approach had an unfavorable impact on the prediction accuracy.

**Figure 6.**
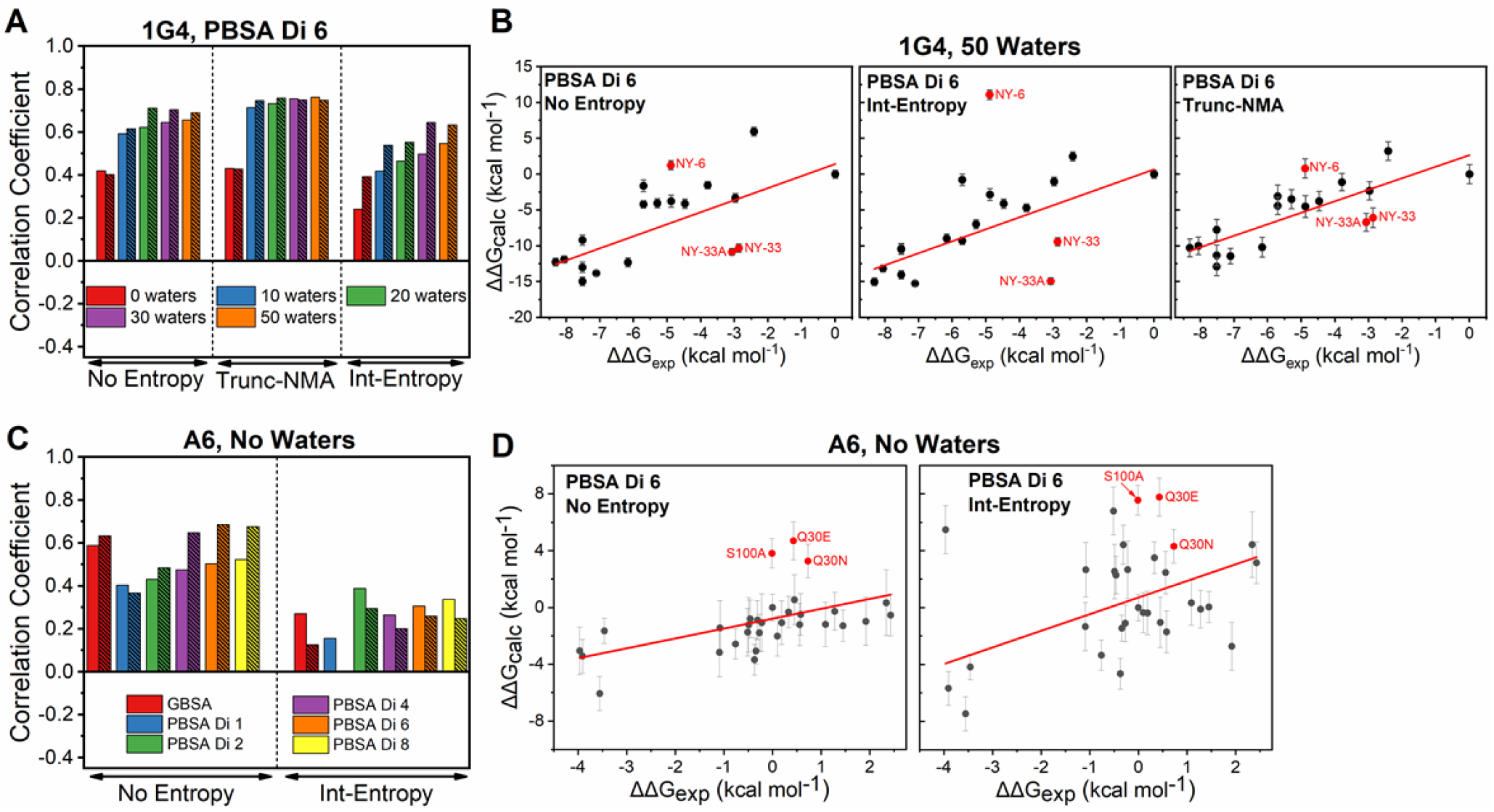
Impact of Solute Entropy Corrections on our MMPB/GBSA Calculations. **A**. Spearman’s rank (rs, unhashed bars) and Pearson’s R (rp, hashed bars) values determined for MMPB/GBSA calculations on the 1G4 test set with 𝜖𝑖𝑛𝑡 set to 6. Results are presented using a variable number of waters without any entropy corrections included as well as with the Trunc-NMA and Int-Entropy approaches. **B**. Exemplar scatter plots for the 1G4 test set with the PBSA approach (with 𝜖𝑖𝑛𝑡 set to 6) including 50 explicit water molecules. Panels compare no entropy corrections (left), with Int-Entropy corrections (middle) and with Trunc-NMA corrections (right). **C**. Impact of the inclusion of the Int-Entropy correction to the A6 data set, with the rs and rp values colored as in **A**. All results are without any explicit water molecules included. **D**. Exemplar scatter plots for the A6 test set with the PBSA approach (with 𝜖𝑖𝑛𝑡 set to 6) and no explicit water molecules. Panels compare no entropy corrections (left), to with Int-Entropy corrections (right). More complete results, including comparing the effect of removing outliers, are provided in **Figure S11**.

Whilst the Int-Entropy has been successfully applied to several small-molecule MMPB/GBSA studies, its application to PPIs has proven more challenging.^72^ This is largely a consequence of the large binding interfaces (TCR-pHLA buried surface areas tend to be ∼2000-2500 Å^2^) which give rise to a correspondingly large amount of variance in the obtained per-frame interaction energies. Thus, without exhaustive sampling, this approach can lead to non-converged and abnormally high entropy corrections.^72^ Further, Ekberg and Ryde have recently argued this method to be intractable for simulations with a large variance in energy, such as the large systems studied here.^73^ One solution to this problem is to perform MMPBSA calculations using an 𝜖_𝑖𝑛𝑡_ value larger than the default of 1, which notably reduces the variance of the interaction energies obtained, ultimately leading to converged entropy estimates within reasonable simulation times.^16^ We indeed observed this behavior with our Int-Entropy corrections for the different MMPBSA methods used in this study, in which only 𝜖_𝑖𝑛𝑡_ values between 2 and 8 showed gaussian-like distributions of the gas phase interaction energy (**Figure S10**). Regardless, the error/noise associated with the calculation was observed to worsen the prediction accuracy for both test sets. We note that when the Int-Entropy method was first introduced by Duan *et al.,* interaction energies were computed using 100,000 snapshots from a single 2 ns long simulation.^44^ In contrast, here we extracted significantly fewer snapshots (7500 frames taken from 25, 3 ns long replicas) and our snapshots were significantly less correlated with one another (frames were taken every 0.02 ps by Duan et al.^44^ instead of every 10 ps here). Whilst some more recent attempts have successfully applied the Int-Entropy approach using notably fewer simulation frames than those used in the original study,^54, 71, 72^ collecting a much larger number of frames to assist with convergence would be significantly more resource intensive, both in terms of the additional MMPB/GBSA calculations needed and the additional storage requirements for the simulations.

We performed Trunc-NMA calculations on only the 1G4 test set (**Figures 6A+B** and **S9**) and obtained no notable change in the prediction accuracy when applying the method to MMPB/GBSA calculations without explicit water molecules. However, the combination of the explicit waters and Trunc-NMA corrections gave rise to a better prediction quality both when including and excluding the aforementioned three outliers. We further note that the improved prediction accuracy associated with Trunc-NMA corrections is not sensitive to the number of explicit water molecules included in the MMPBSA calculation (**Figure 6A**; similar as observed without applying entropy corrections, **Figure 4**).

Whilst we did not evaluate the non-truncated form of NMA, previous studies have clearly shown the beneficial effects of using a truncated system on both the errors obtained and computational efficiency.^45, 68^ Given the size of a standard TCR-pHLA complex (∼800-900 residues), the Trunc-NMA approach used here would be significantly more efficient than standard NMA. For the 1G4 data set composed of many multi-point mutations, the combination of Trunc-NMA and explicit water molecules was beneficial according to all three metrics we evaluated (rs and rp in **Figure 6A+B** and the MAD in **Table S7**). Further, we observed the prediction quality to be highly insensitive to the number of explicit water molecules included in the MMPBSA calculation.

### How many replicas are required for reproducible MMPB/GBSA calculations?

The results presented so far have shown a clear benefit of the use of a 𝜖_𝑖𝑛𝑡_ value ≥ 4 for MMPBSA calculations, both in terms of improving the prediction quality and in reducing the magnitude of the errors obtained. Further, beneficial effects were also observed for the 1G4 test set when both explicit water molecules and entropy corrections were applied. However, these methods are likely to increase the noise associated with the predictions. It is therefore important to assess how many replicas may be required for reproducible results with the different approaches performed in this study. We used ‘bootstrapping’ to do this: a statistical method that involves “resampling with replacement”, meaning that from a set of N observables (in our case N is the 25 replicas performed for each complex) a large number of bootstrap “resamples” are constructed by randomly removing or duplicating the individual observations.

These bootstrap resamples are then used to recalculate the correlation coefficients many times in order to obtain confidence intervals in the calculated correlation coefficients. For both test sets, we generated 1,000,000 bootstrap resamples of ΔΔ*G*calc for several different MMPB/GBSA protocols used here. We then evaluated the impact of using a reduced number of replicas on the confidence intervals of the Spearman’s rank (**Figure 7**) and Pearson’s R (**Figure S12**). We observed very similar behavior for both measures, so only Spearman’s rank (**Figure 7**) is discussed below. We note that each average correlation coefficient value in **Figure 7** it is not an informative metric for determining a suitable sample size, as it is determined from (up to) a million randomly selected resamples. Instead, the size of the confidence intervals (and how much they are reduced with an increasing number of replicas) is a measure of how reproducible the results would be (for a given number of replicas).

**Figure 7.**
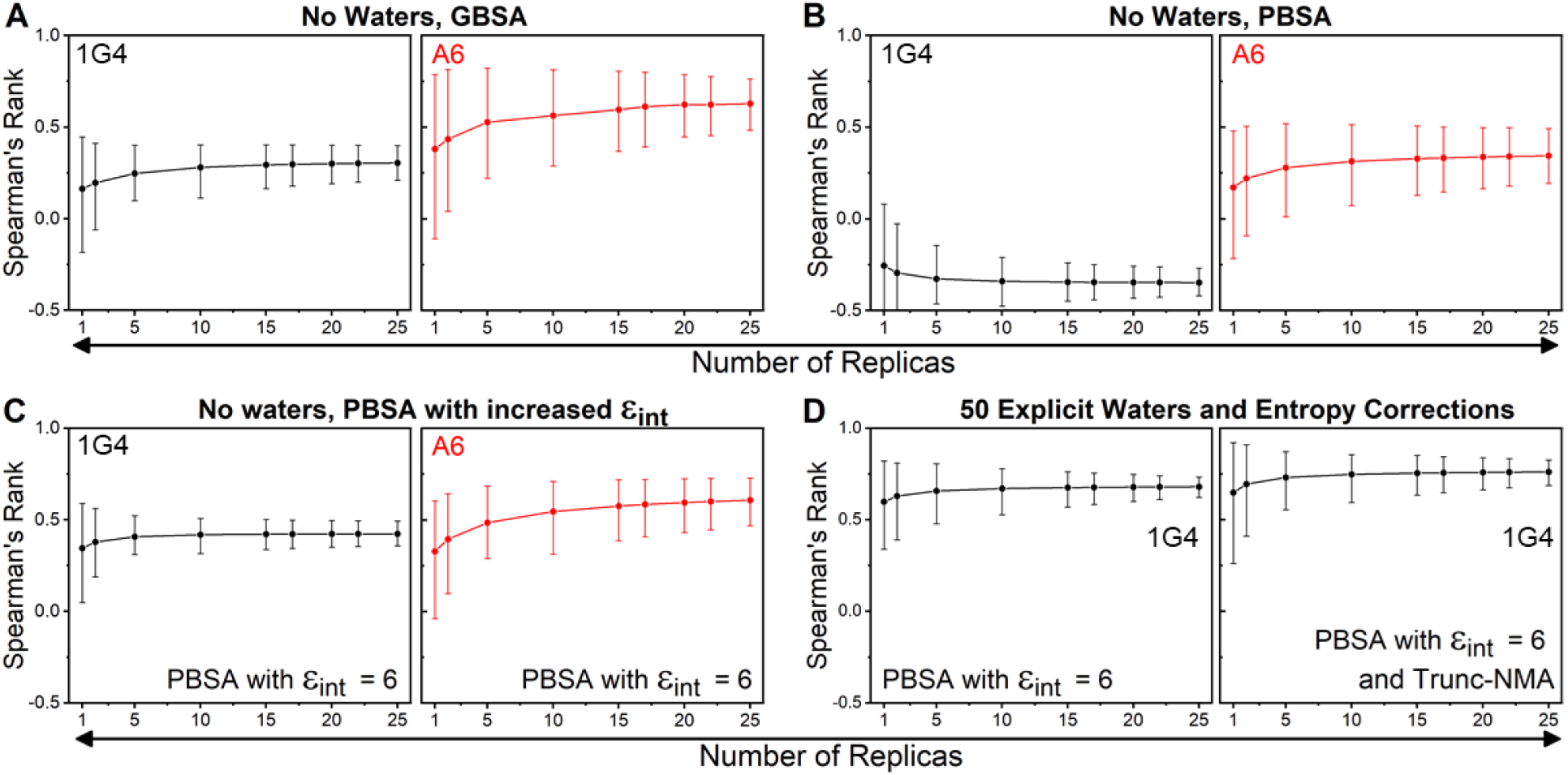
Bootstrapping to assess the impact of using differing numbers of replicas to obtain Spearman’s Rank for some of the protocols evaluated in this study. Panels **A+B** focus on the GBSA and PBSA approaches with no explicit waters included. Panel **C** focusses on the PBSA method with 𝜖𝑖𝑛𝑡 set to 6. Panel **D** focusses on the PBSA method (𝜖𝑖𝑛𝑡 set to 6) with 50 explicit waters molecules included with and without the Trunc-NMA correction applied. Measurements with the 1G4 and A6 test sets are colored black and red respectively. In each panel, the average of the 1 million bootstrap resamples are used to calculate Spearman’s Rank when using a differing number of replicas, with the error bars depicting the 95% confidence intervals. The complete data is used in all cases (*i.e.,* the outliers discussed above are included). Equivalent results with the Pearson’s R metric are provided in **Figure S12.**

Focusing first on the 1G4 test set (**Figure 7A**-**D**), there appears to be little benefit for performing more than 15 replicas for the MMGBSA approach, whilst for the MMPBSA simulations with 𝜖_𝑖𝑛𝑡_ set to 6, one could argue that as few as 5 replicas may be sufficient, considering the additional computational cost if more replicas are used. This is also true when explicit water molecules are included and/or Trunc-NMA entropy corrections applied: 5-10 replicas are sufficient to converge the prediction estimates. Comparison of the A6 and 1G4 test sets shows that the A6 test set is generally noisier for each comparable method (**Figure 7A-C**).

This is likely in part due to the reduced experimental affinity range in the data set as well as the comparably lower quality of the WT crystal structure (resolutions of 1.9 Å vs 2.6 Å for 1G4^31^ and A6^32^, respectively). For the A6 test set, a larger number of replicas may therefore be optimal as compared to the 1G4 TCR, in terms of the balance between accuracy and computing cost. Regardless, for both test sets a maximum of 15 replicas would appear to be sufficient when using the optimal parameters previously described.

## Conclusions

Here, we evaluated MMPB/GBSA binding affinity calculation protocols for two contrasting TCR-pHLA test sets: 1G4, with 3-14 mutations across a number of CDR loops^26, 46^ and A6, with primarily single mutations on a single CDR loop (CDR3β).^29, 33^ Although there is no single protocol that is highly suitable for both sets, there are general lessons to be learned and specific recommendations for the application of MMPB/GBSA to TCR-pHLA complexes that can be made based on our results.

First, an increased value (between 4-8) of 𝜖_𝑖𝑛𝑡_ is strongly recommended for MMPBSA calculations. This should improve prediction quality *and* fewer simulations are required per complex (e.g. 5-10 simulations of 4 ns, see **Figure 7**). Second, there is a divergence in the optimal protocol between our two test sets regarding inclusion of explicit water molecules: For the 1G4 set this may improve prediction accuracy, whereas for the A6 set this led to reduced accuracy (due to additional errors/noise). Third, using truncated NMA entropy corrections improved prediction accuracy when variants had significantly altered H-bonding across the interface (thus resolving significant outliers), whereas using ’interaction entropy’ corrections is not suitable.

Overall, we recommend the following for TCR-pHLA relative binding affinity prediction with MMPBSA: (1) Use an internal dielectric constant of ∼6; (2) A truncated NMA based entropy correction should be applied when mutations cause significant changes in the TCR-pHLA hydrogen bonding network; (3) Inclusion of explicit water molecules at the interface should be done with caution, as it can increase noise. When computational efficiency is important, MMGBSA could be considered for TCR variants with few mutations.

Finally, our bootstrapping analysis demonstrated that for MMPBSA as few as 5 replicas (20 ns MD in total) can be sufficient to obtain reproducible results. Thus, in a practical context, one could envisage evaluating candidate variants initially using 5 replicas, followed by completing a total of 10-15 replicas for promising variants for increased accuracy.

Computational methods that allow for the accurate ranking of TCR-pHLA binding affinities and those of PPIs more generally have obvious utility in computational drug discovery. Whilst we intended to find a general approach, our results demonstrated the need for two somewhat different approaches for accurate and reliable ranking of TCR-pHLA binding affinities: one for ranking TCR variants with multiple mutations (>4), and one with few mutations. We believe the MMPB/GBSA approach outlined here has promise as a medium throughput screening tool to select and rank candidate TCR variants for experimental testing.

## Author contributions

RMC and MWvdK designed the study. RMC performed the simulations and analysis. RMC produced the first draft manuscript which was edited through contributions from all authors. All authors discussed and interpreted data. All authors have given approval to the final version of the manuscript.

## Acknowledgments

The authors would like to thank Dr. Dimas Suarez (Univ. of Oviedo, Spain) for assistance in modifying the Nmode program to enable the running of Trunc-NMA calculations.

## Abbreviations

CDR: Complementarity-determining region
Int-Entropy: interaction entropy
𝜖_𝑖𝑛𝑡_: internal dielectric constant
MD: molecular dynamics
MMPB/GBSA: Molecular Mechanics Poisson-Boltzmann/generalized Born surface area
pHLA: peptide-human leukocyte antigen
PBA: polar buried area
PPI: protein-protein interaction
*g*(***r***): radial distribution function
TCR: T-cell receptor
Trunc-NMA: truncated normal mode analysis

## Data availability

All starting structures for simulation (Amber topology files and coordinates), example input files for MD simulations and MMPB/GBSA calculations, example analysis scripts and a copy of the modified NMode code used are available at DOI: 10.5281/zenodo.4805388. All further relevant data are within the manuscript and the **Supplementary Information**.

