## Supporting Information for "Reliable *in silico* ranking of engineered therapeutic TCR binding affinities with MMPB/GBSA"

*** Corresponding authors**

### Additional methods

*WT A6-TCR Solvation.* The 3D-Reference Interaction Site Model[1,2] (3D-RISM) was used to predict the density distribution function ($g(\boldsymbol{r})$) for water oxygen atoms across the entire TCR-pHLA binding site of the WT A6 TCR-pHLA structure. For 3D-RISM calculations (performed with AmberTools18), the Kovalenko-Hirata (KH) closure method[3,4] was used, with all other settings kept as default. Following this, Placevent[5] was used to solvate the entire TCR-pHLA complex, by solvating the entire complex with waters molecules up to 5 Å away from any protein atom. Multiple cut-off $g(\boldsymbol{r})$ values (point at which additional waters with smaller $g(\boldsymbol{r})$ values are no longer added) were tested as the default value of 1.5 Å resulted in stability issues (due to "vacuum bubbles") during NPT simulations (due to too much space between solvent atoms in the initial structure). We found that appropriate solvation was achieved with a $g(\boldsymbol{r})$ cut-off of 1.1 Å and following this we solvated the protein and these water molecules in a water box of size 7 Å (away from both any protein and 3D-RISM/Placevent water molecule). This distance of 7 Å was chosen as it gave a slightly bigger box size (in all dimensions) to the box size that would be generated when solvating just the protein in a 10 Å octahedral water box. (This value is likely to be somewhat system dependent.)

*Structure equilibration procedure.* The following procedure was used to prepare all systems simulated for production MD simulations in the NPT ensemble at 298 K and 1 atm. All dynamics steps applied the SHAKE algorithm to constrain all bonds containing a hydrogen atom. Replica simulations were initiated from the second heating step of the following protocol (with each replica assigned different random velocity vectors at this stage). Simulations performed in the NVT ensemble used Langevin temperature control (with a collision frequency of 1 $ps^{-1}$) and used a simulation timestep of 1 fs. Simulations performed in the NPT ensemble used Langevin temperature control (collision frequency of 1 $ps^{-1}$) and a Berendsen barostat (1 ps pressure relaxation time).

The equilibration protocol is as follows: First, hydrogens atoms and solvent molecules were energy minimized (using 500 steps of steepest descent followed by 500 steps of conjugate gradient minimization). To prevent the movement of non-hydrogen and non-solvent atoms during the minimization, 10 kcal $mol^{-1}$ $Å^{-1}$ positional restraints were used to keep all heavy atoms fixed. Then the solvent was heated rapidly from 50 K to 298 K (NVT ensemble, 1 fs timestep) over the course of 200 ps, with the previously described restraints still maintained. The positional restraints were then replaced with 5 kcal $mol^{-1}$ $Å^{-1}$ positional restraints on only the Cα carbon atoms of each residue and subjected to another round of energy minimization

(500 steps of steepest descent followed by 500 steps of conjugate gradient). Retaining these positional restraints, the system was heated from 25 K to 298 K over the course of 50 ps (NVT ensemble, 1 fs time step). Simulations were then performed in the NPT ensemble (1 atm, 298 K, 2 fs time step) by first gradually reducing the 5 kcal $mol^{-1}$ $Å^{-1}$ Cα carbon restraints over the course of 50 ps. This was done by reducing the restraint weight by 1 kcal $mol^{-1}$ $Å^{-1}$ every 10 ps. A final 1 ns long NVT MD simulation with no restraints placed on the system was then performed, with the final structure produced after this run, used as the starting point for production MD simulations.

*Truncated Normal Mode Analysis.* Truncated normal mode analysis calculations (Trunc-NMA) were performed only on the 1G4 TCR-pHLA complexes studied in this manuscript. Residues in the pHLA that were retained were HLA residues: 5-26, 33-47, 54-101, 112-118, 123-133, 139-174, and peptide residues: 1-9. Residues in the TCR that were retained were TCRα-chain: 1-5, 23-36, 44-73, 89-105 and TCRβ-chain: 1-4, 22-34, 42-75, 89-104. Residues that were restrained during energy minimization and vibrational frequency calculations were: HLA residues: 6, 8, 10, 15-16, 23, 25-26, 33-35, 47, 54, 85-88, 93-94, 96, 98, 100-101, 112-113, 115, 117-118, 123, 125, 127-129, 139-141, 169 and172-174; TCRα-chain residues: 4-5, 23-24, 26, 35, 45-48, 60-65, 67, 67-73, 89-91, 105; and TCRβ-chain residues: 1, 3, 22, 42-43, 59, 61-64, 74-75, 89-90, 104. The "closest" command in the Ambertools18 program CPPTRAJ[6] was used to retain 1000 binding site water molecules in each frame. The 1000 closest water molecules (water position determined using the oxygen atom) to the Cα of any of central peptide residues (residues 4, 5, 6 and 7) were used for this closest water molecules calculation. We used a modified version of the Nmode program from AmberTools14, which allows for the use of the "ibelly" parameter which fixes selected atoms in place, meaning they have no (direct) contribution to the vibrational frequency calculation. The modified Nmode code (along with scripts to setup and analyse the obtained results) can be provided upon request to the corresponding authors.

*Inclusion of Explicit Water Molecules for MMPB/GBSA Calculations.* MMPB/GBSA calculations performed with explicit solvent (either 10, 20, 30 or 50 water molecules) were selected for using the "closest" command with CPPTRAJ.[6] Water molecules were included in the complex and receptor calculations, and receptor residues at the binding site were used to select the $x$ closest water molecules. Specifically, for WT 1G4, HLA residues: 19, 62, 65-66, 68, 69, 71-73, 75, 76, 150, 151, 154, 155, 158, 159, 163, 167 and peptide residues: 4-8 were used. For WT A6, HLA residues: 62, 65, 66, 69, 72-73, 150, 155, 158, 163, 166 and peptide residues: 1-7 were used. The *x* closest water oxygen atom distances to any nitrogen or oxygen atom on the defined receptor residues were then used. The following selection expression was used in CPPTRAJ: “(:receptor_residue_selection)&(@O*|@N*)”.

### Additional figures

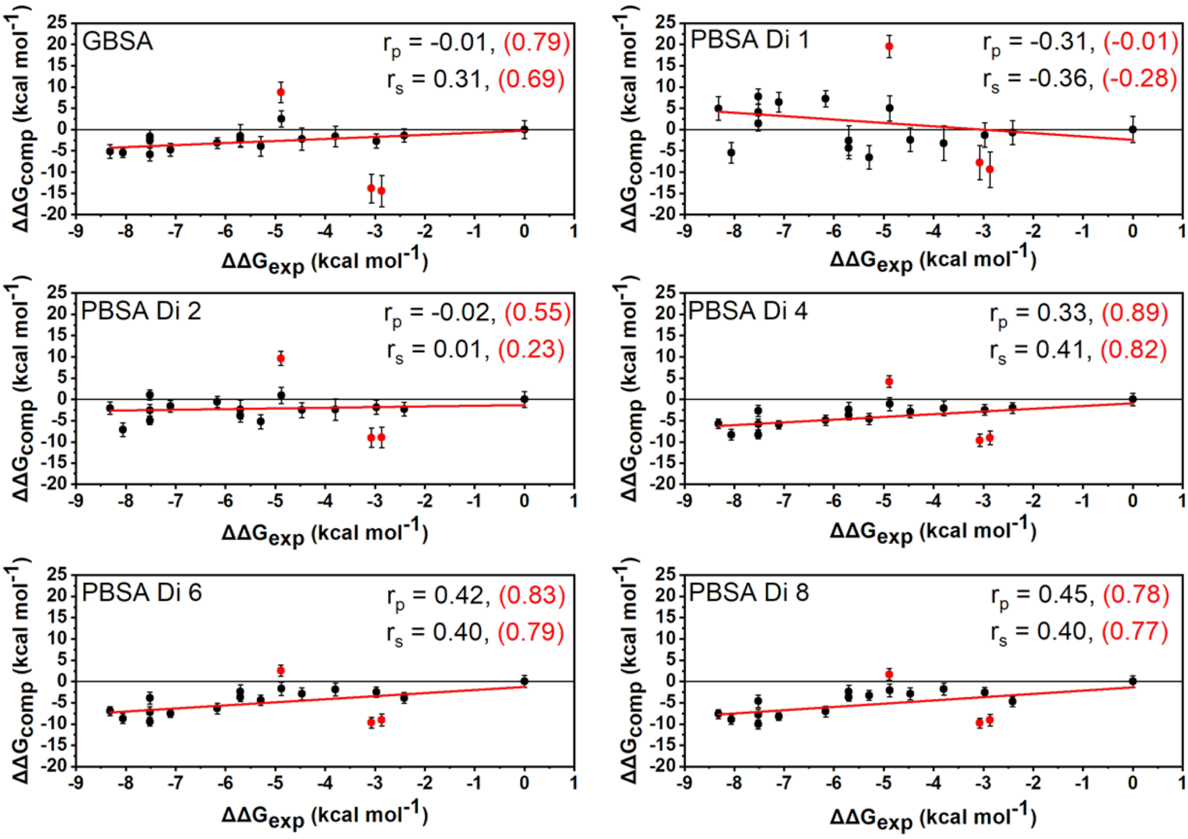

**Figure S1.** Computational *versus* experimental $\Delta\Delta G$ values obtained for the 1G4 set of TCRs. For each panel, the MMGB/PBSA methods used are indicated in the top right. The red line corresponds to a linear fit of all the data. The three variants colored in red correspond to the outliers (NY-6, NY-33 and NY-33A) discussed in the main text. The Pearson's r value ($r_p$) and Spearman's rank ($r_s$) are provided for the full set of data (in black) and without the three identified outliers discussed in the main text (in red and in parenthesis).

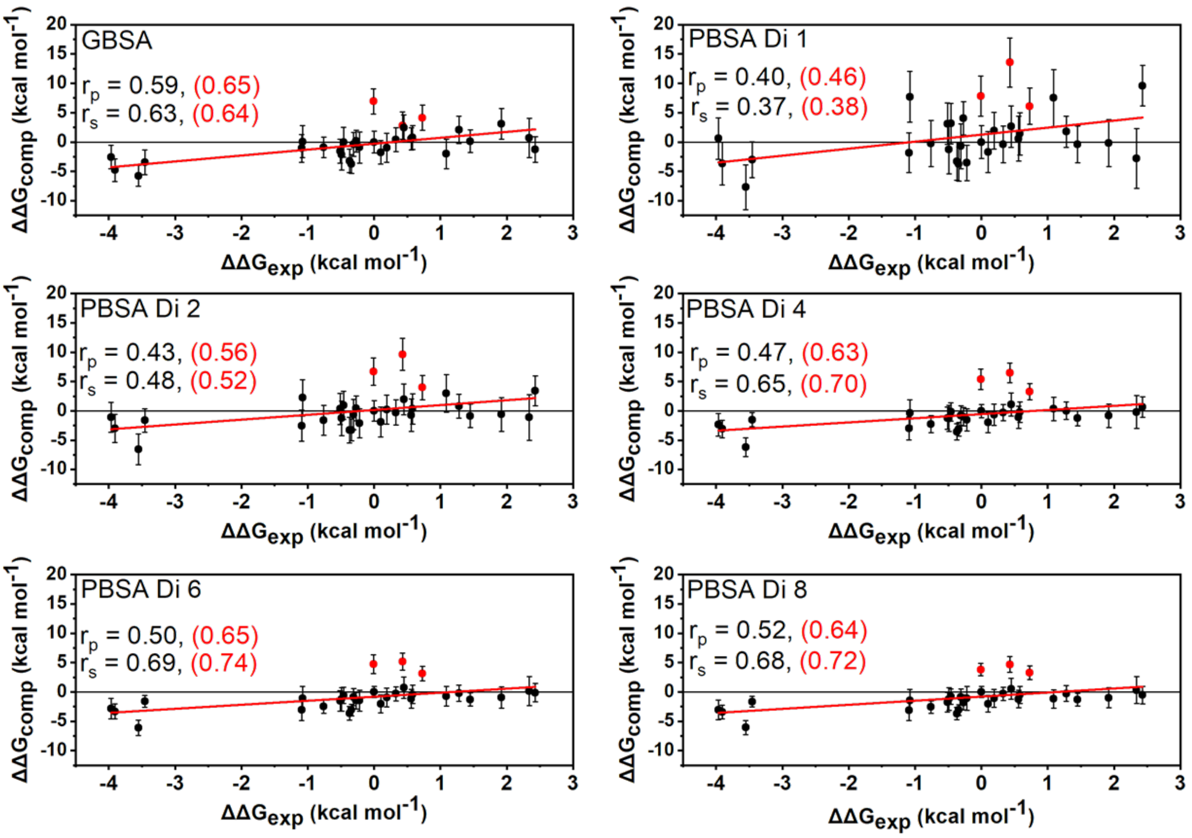

**Figure S2.** Computational *versus* experimental $\Delta\Delta G$ values obtained for the A6 set of TCRs. For each panel, the MMGB/PBSA methods used are indicated in the top right. The red line corresponds to a linear fit of all the data. The Pearson's r value ($r_p$) and Spearman's rank ($r_s$) for each method applied are provided for the full set of data (in black) and without the three identified outliers discussed in the main text (in red and in parenthesis).

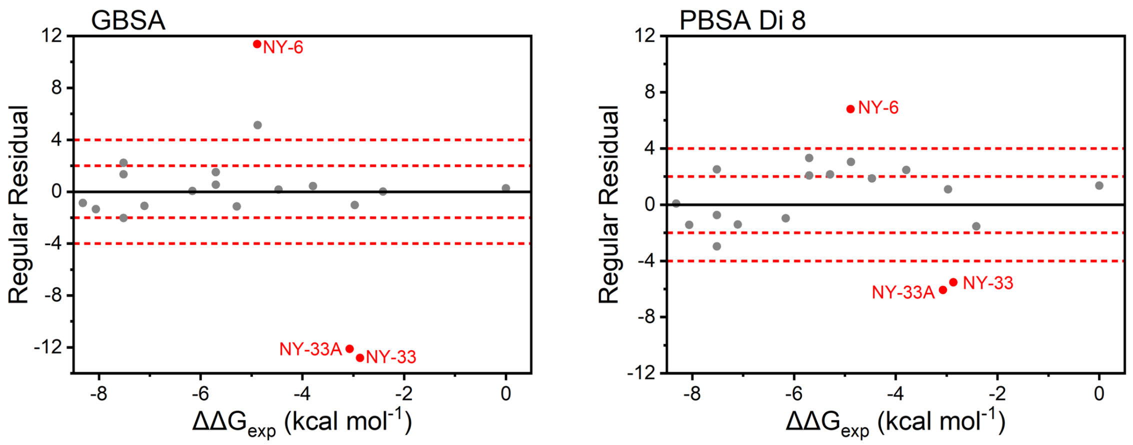

**Figure S3**. Residual plots of the 1G4 TCR test set for MMGBSA calculations (left panel) and MMPBSA calculations with the internal dielectric constant set to 8 (right panel), in the absence of any explicit water molecules or solute entropy corrections. (We note that results for MMPBSA calculations with the internal dielectric set to either 4 or 6 were highly similar to results shown above for a dielectric constant of 8.) The three TCR variants described as outliers in the main text are colored red.

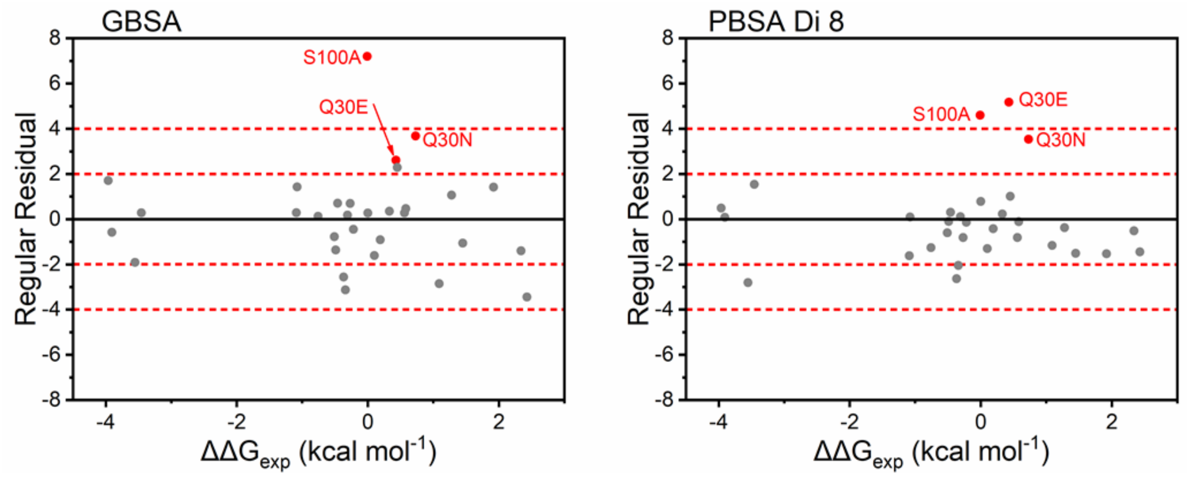

**Figure S4**. Residual plots of the A6 TCR test set for MMGBSA calculations (left panel) and MMPBSA calculations with the internal dielectric constant set to 8 (right panel), in the absence of any explicit water molecules or solute entropy corrections. (We note that results for MMPBSA calculations with the internal dielectric set to either 4 or 6 were highly similar to results shown above for a dielectric constant of 8.) The three TCR variants described as outliers in the main text are colored red as their MMPBSA calculations had residuals greater than 3 for the MMPBSA calculations.

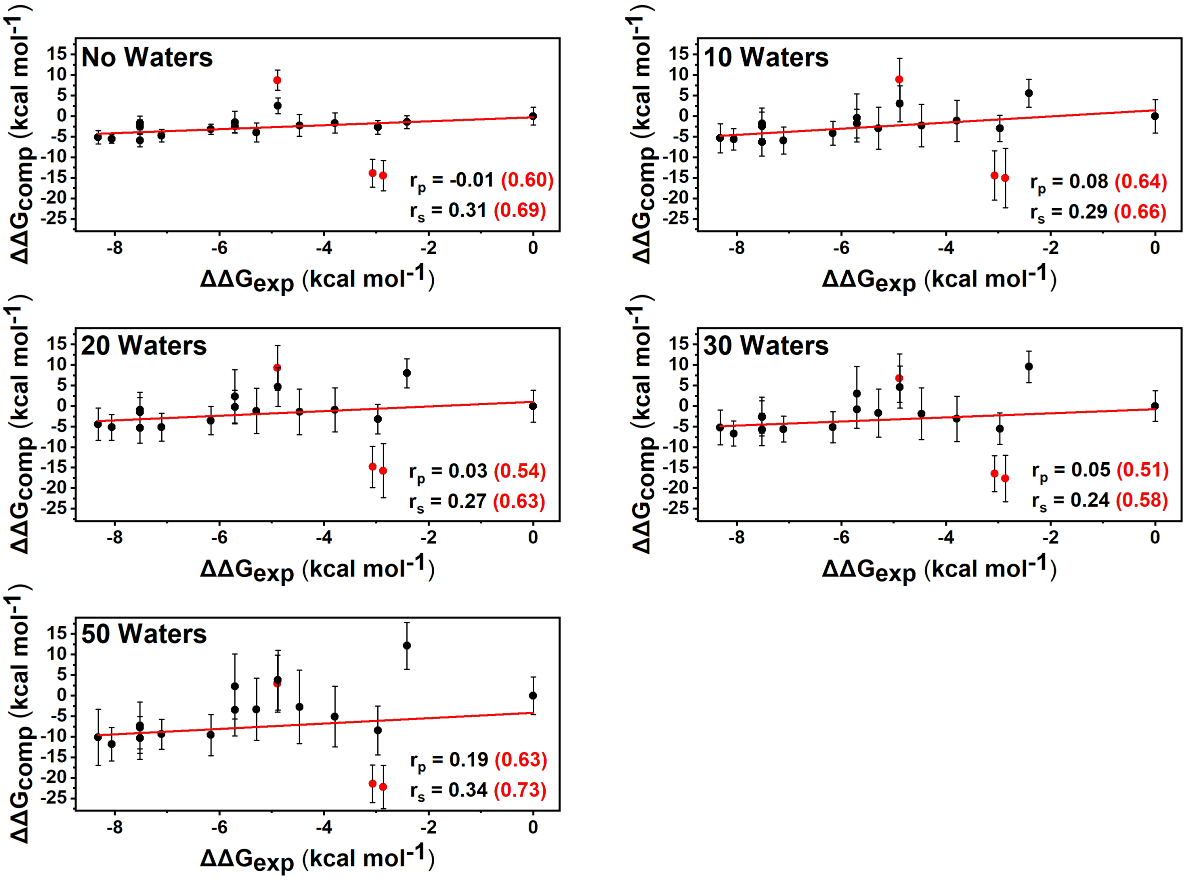

**Figure S5.** Computational *versus* experimental Δ$\Delta G$ values obtained for the 1G4 set of TCRs using the MMGBSA method with different amounts of explicit waters included. The number of waters used for each calculation is stated in the top left corner of each plot. The red line corresponds to a linear fit of all the data. The three variants colored in red correspond to the outliers (NY-6, NY-33 and NY-33A) discussed in the main text. The Pearson's r value ($r_p$) and Spearman's rank ($r_s$) are provided for the full set of data (in black) and without the three identified outliers discussed in the main text (in red and in parenthesis).

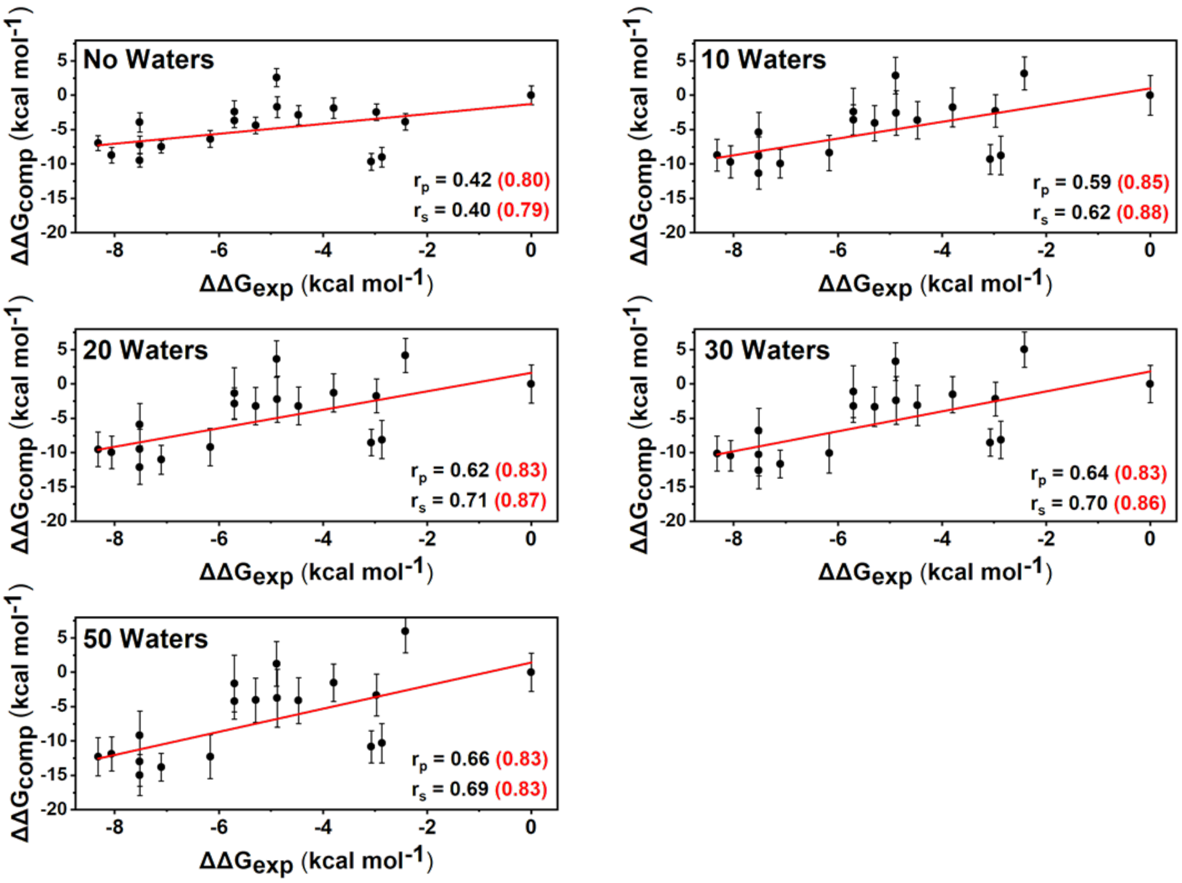

**Figure S6.** Computational *versus* experimental $\Delta\Delta G$ values obtained for the 1G4 set of TCRs using the MMPBSA method with the internal dielectric constant set to 6 and a varied number of waters included. The number of waters used for each calculation is stated in the top left corner of each plot. The red line corresponds to a linear fit of all the data. The three variants colored in red correspond to the outliers (NY-6, NY-33 and NY-33A) discussed in the main text. The Pearson's r value ($r_p$) and Spearman's rank ($r_s$) are provided for the full set of data (in black) and without the three identified outliers discussed in the main text (in red and in parenthesis).

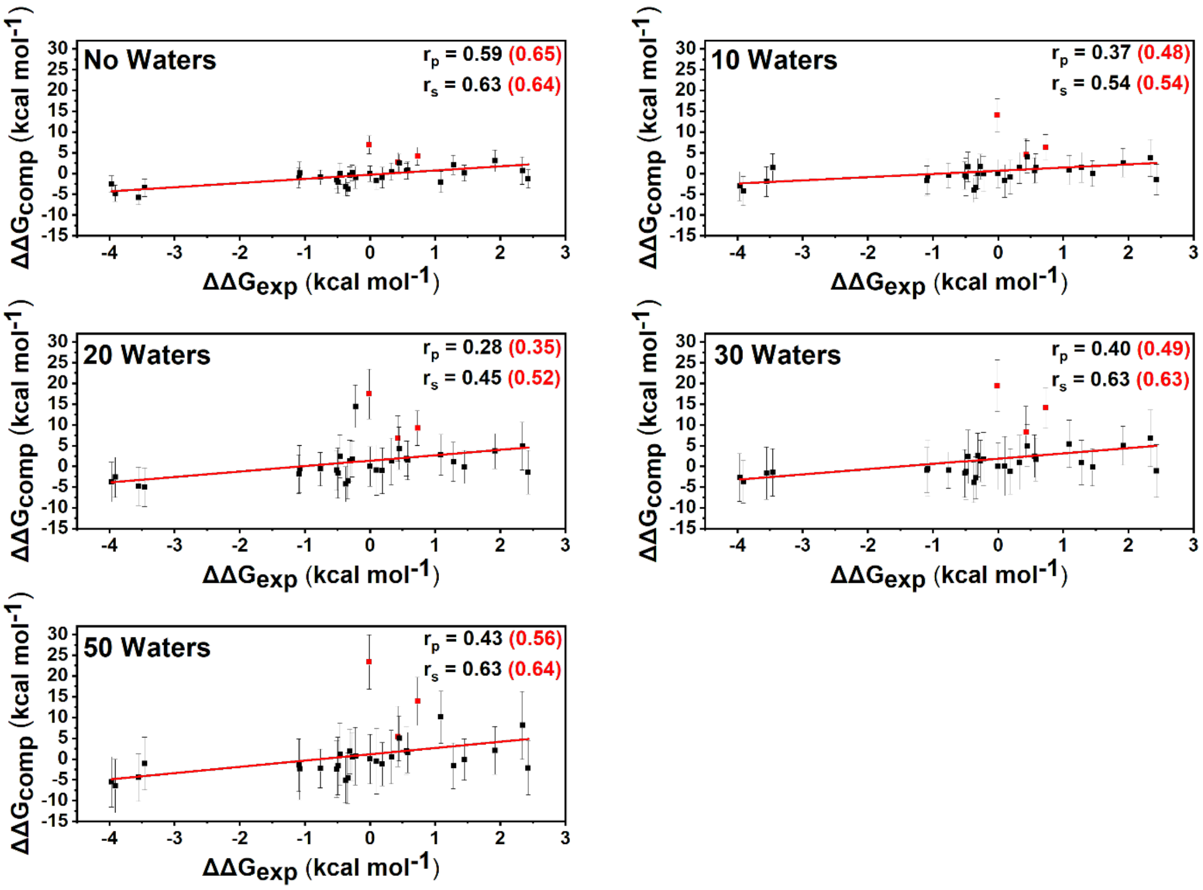

**Figure S7.** Computational *versus* experimental ΔΔ*G* values obtained for the A6 set of TCRs using the MMGBSA method and a variable amount of explicit water molecules. The number of waters used for each calculation is stated in the top left corner of each plot. The red line corresponds to a linear fit of all the data. The three variants colored in red correspond to the outliers (point mutants Q30E, Q30N and S100A) discussed in the main text. The Pearson's r value ($r_p$) and Spearman's rank ($r_s$) are provided for the full set of data (in black) and without the three identified outliers discussed in the main text (in red and in parenthesis).

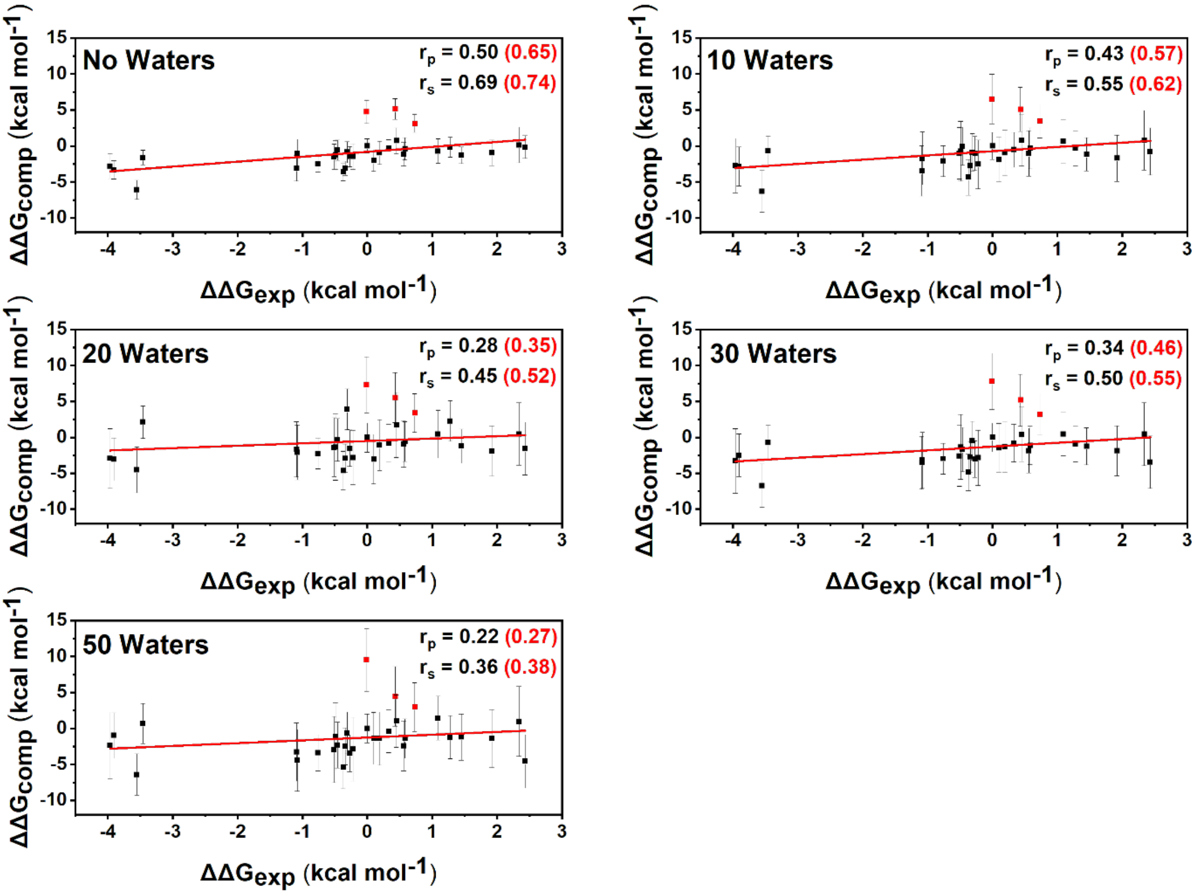

**Figure S8.** Computational *versus* experimental $\Delta\Delta G$ values obtained for the A6 set of TCRs using the MMPBSA method with the internal dielectric constant set to 6 and with a varied number of waters included. The number of waters used for each calculation is stated in the top left corner of each plot. The red line corresponds to a linear fit of all the data. The three variants colored in red correspond to the outliers (point mutants Q30E, Q30N and S100A) discussed in the main text. The Pearson's r value ($r_p$) and Spearman's rank ($r_s$) are provided for the full set of data (in black) and without the three identified outliers discussed in the main text (in red and in parenthesis).

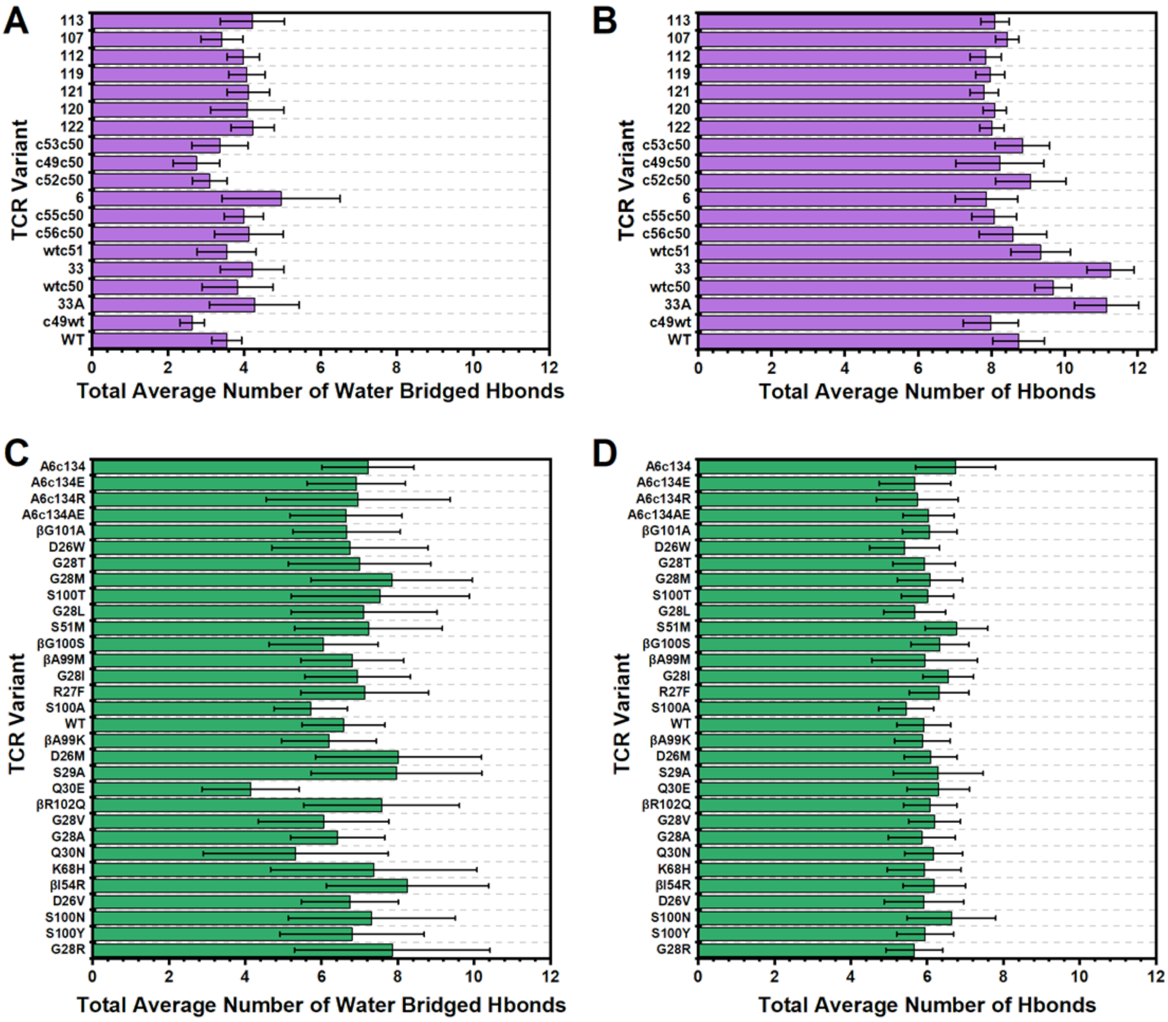

**Figure S9.** Total average number of water bridged and solute-solute hydrogen bonds (Hbonds) formed between the TCR and pHLA for each TCR variant simulated. Results for both the 1G4 (**A**+**B**) and A6 (**C**+**D**) test sets are shown. Error bars are the standard deviation from the 25 replicas performed per complex. Water bridged Hbonds for the 1G4 and A6 TCRs are shown in panels **A** and **C** respectively, whilst solute-solute Hbonds are shown in panels **B** and **D** respectively. For the A6 TCRs (panels **C**+**D**), single point mutations preceded by the letter "β", mean the mutation is located on the TCRβ chain (with all others therefore located on TCRα chain). A hydrogen bond is defined as having a donor-acceptor distance less than 3.5 Å, and a donor-hydrogen-acceptor angle within 180 ± 45°. For each graph, the TCR variants are ordered according to their affinity (highest affinity at the top, lowest affinity at the bottom).

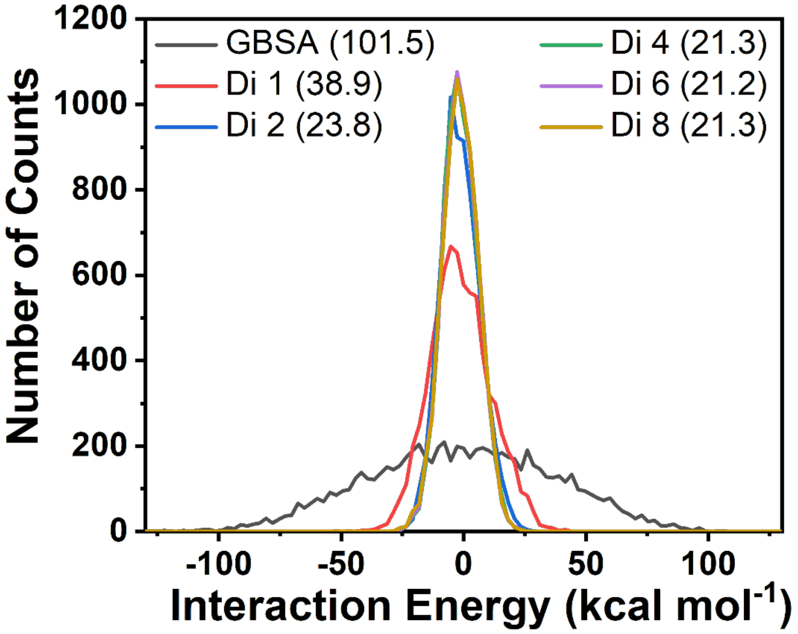

**Figure S10.** Histograms of the fluctuation of gas phase interaction energy. Histograms were obtained by solving **Equation 4** in the main text for the WT-1G4 TCR-pHLA complex using the six different protocols evaluated in this study (performed with no explicit water molecules included). The value in brackets corresponds to the entropy correction estimate i.e. $-T\Delta S$ (in kcal mol$^{-1}$) obtained from each method when using the interaction entropy (Int-Entropy) approach (**Equation 5** in the main text).

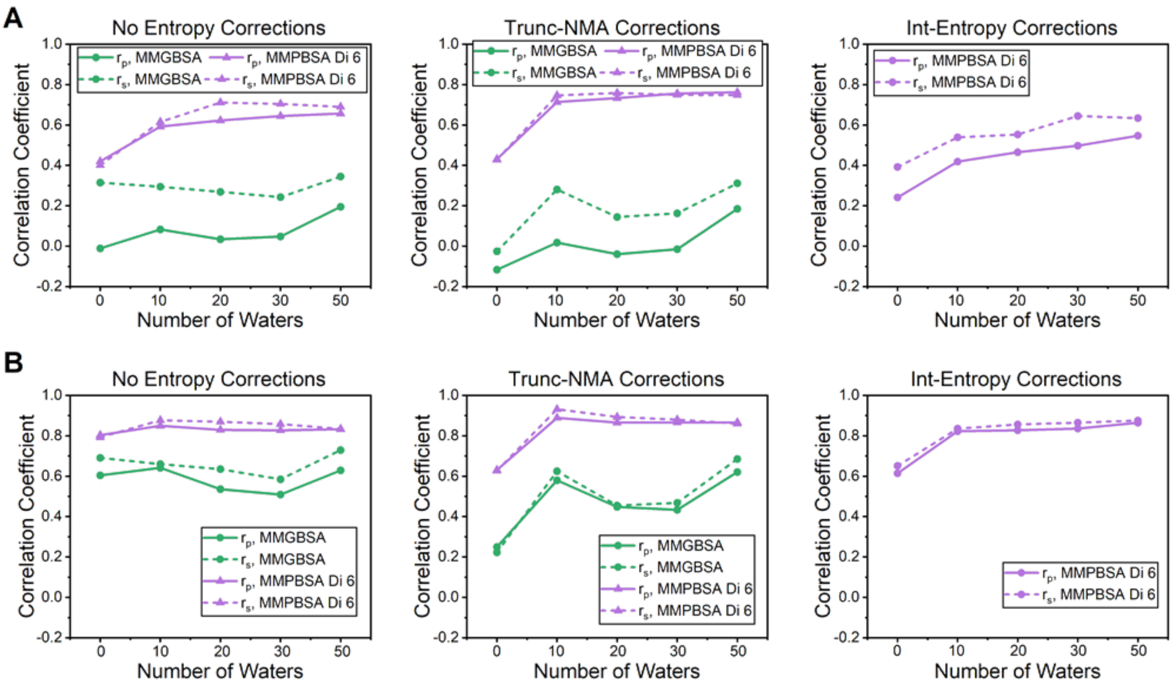

**Figure S11.** Impact of entropy corrections to the 1G4 test set as determined by the Pearson's R ($r_p$) and Spearman's rank ($r_s$). The three panels in section (**A**) contain the results: No entropy corrections (left) with truncated normal mode analysis (Trunc-NMA) corrections (middle) and Interaction Entropy (Int-Entropy) corrections with all data included (*i.e.* outliers described in the main text are included). The three panels in section (**B**) show the same data but with the three outliers described in the main text excluded from the calculations.

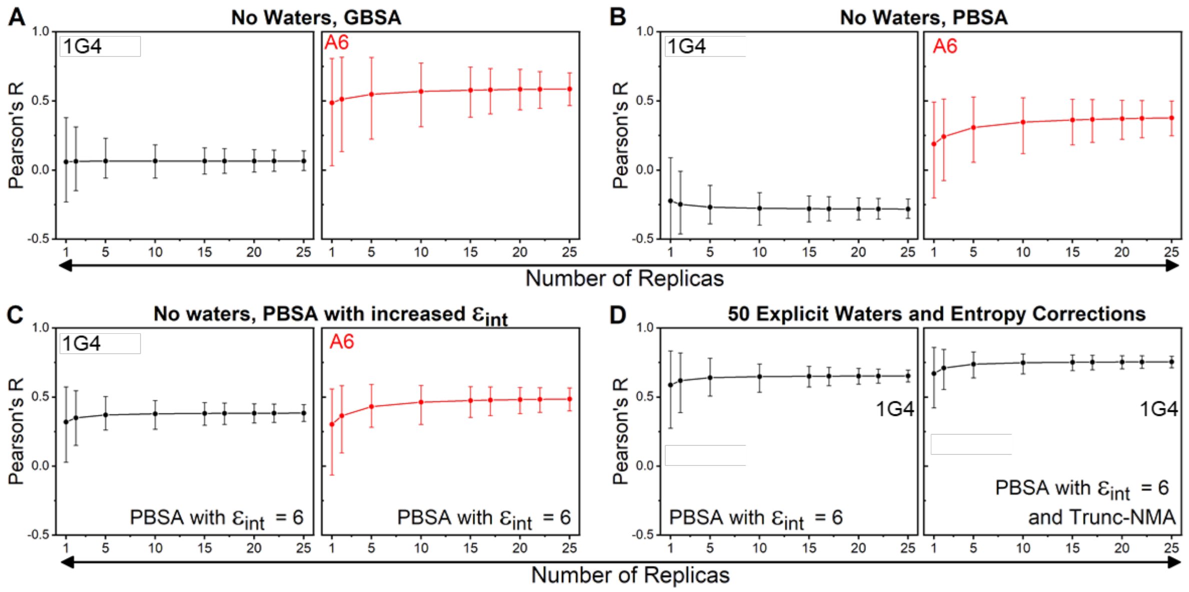

**Figure S12.** Bootstrapping to assess the impact of the number of replicas on the Pearson's R value for some of the protocols evaluated in this study. Panels **A**+**B** focus on the GBSA and PBSA approaches with no explicit waters included. Panel **C** focusses on the PBSA method with $\epsilon_{int}$ set to 6. Panel **D** focusses on the PBSA method ($\epsilon_{int}$ set to 6) with 50 explicit waters molecules included with and without the Trunc-NMA correction applied. Measurements with the 1G4 and A6 test sets are colored black and red respectively. In each panel, 1 million bootstrap resamples are used to calculate Spearman's Rank when using a differing number of replicas. Error bars shown are the 95% confidence intervals. The complete data is used in all cases (i.e. the outliers discussed above are included). Equivalent results for the Spearman's rank metric are provided in the main text (**Figure 7**).

### Supporting tables

**Table S1.** Histidine tautomerization states used for 1G4 and A6 TCR-pHLA complex MD simulations. Numbering is consistent with PDB ID 2BNR[7] for 1G4 and PDB ID 1AO7[8] for A6 simulations.

| TCR-pHLA System | HID[a] Tautomers | HIE[b] Tautomers |
|---|---|---|
| 1G4 | HLA: 3, 70, 74, 93, 114, 192, 260.<br>β2m: 51.<br>Peptide: N/A.<br>TCRα: N/A.<br>TCRβ: 152. | HLA: 145, 151, 188, 191, 197, 263.<br>β2m: 13, 31, 84.<br>Peptide: N/A.<br>TCRα: 113.<br>TCRβ: 28, 46, 135, 165, 205. |
| A6 | HLA: 70, 74, 93, 114, 191.<br>β2m: N/A.<br>Peptide: N/A.<br>TCRα: N/A.<br>TCRβ: N/A. | HLA: 3, 145, 151, 188, 192, 197, 260, 263.<br>β2m: 13, 31, 51, 84,<br>Peptide: N/A<br>TCRα: N/A<br>TCRβ: 29, 47, 139, 156, 169, 209. |

[a] HID corresponds to a histidine residue which is singly protonated on its Nδ1 nitrogen.

[b] HIE corresponds to a histidine residue which is singly protonated on their Nε2 nitrogen.

**Table S2.** Experimental TCR-pHLA affinities for the 1G4 TCRs. TCR IDs used correspond to the same as those in the original paper indicated.

| TCR ID | $K_D$ (nM) | $\Delta G_{bind}$ (kcal mol$^{-1}$) | $\Delta\Delta G_{bind}$ (kcal mol$^{-1}$) | Reference |
|---|---|---|---|---|
| **Data set 1[a]** | | | | |
| **WT** | 32,000 | -6.14 | 0 | Li *et al.*[9] |
| **6** | 8.4 | -11.03 | -4.89 | Li *et al.*[9] |
| **33** | 180 | -9.21 | -3.07 | Li *et al.*[9] |
| **33A** | 254 | -9.01 | -2.87 | Li *et al.*[9] |
| **107** | 0.04 | -14.20 | -8.06 | Li *et al.*[9] |
| **112** | 0.1 | -13.65 | -7.52 | Li *et al.*[9] |
| **113** | 0.026 | -14.45 | -8.32 | Li *et al.*[9] |
| **119** | 0.1 | -13.65 | -7.52 | Li *et al.*[9] |
| **120** | 0.2 | -13.24 | -7.11 | Li *et al.*[9] |
| **121** | 0.1 | -13.65 | -7.52 | Li *et al.*[9] |
| **122** | 0.98 | -12.30 | -6.16 | Li *et al.*[9] |
| **Data set 2[a]** | | | | |
| **WT** | 15,000 | -6.59 | 0 | Dunn *et al.*[10] |
| **c49wt** | 255 | -9.00 | -2.42 | Dunn *et al.*[10] |
| **c49c50** | 1 | -12.29 | -5.70 | Dunn *et al.*[10] |
| **c52c50** | 2 | -11.88 | -5.29 | Dunn *et al.*[10] |
| **c53c50** | 1 | -12.29 | -5.70 | Dunn *et al.*[10] |
| **c55c50** | 4 | -11.47 | -4.88 | Dunn *et al.*[10] |
| **c56c50** | 8 | -11.06 | -4.47 | Dunn *et al.*[10] |
| **wtc50** | 100 | -9.56 | -2.97 | Dunn *et al.*[10] |
| **wtc51** | 25 | -10.38 | -3.79 | Dunn *et al.*[10] |

[a] For both different studies, $\Delta\Delta G$ values were determined using the WT affinity determined in the same study.

**Table S3.** Experimental TCR point mutation data with TCR-pHLA affinities for the A6 TCR series. TCR IDs used correspond to the same as those in the original paper indicated.

| TCR I.D. | $K_D$ (nM) | $\Delta G_{bind}$ (kcal mol$^{-1}$) | $\Delta\Delta G_{bind}$ (kcal mol$^{-1}$) | Reference |
|---|---|---|---|---|
| **Data set 1 (all single point mutations)**[a] | | | | |
| **WT** | 3200 | -7.50 | 0 | Haidar *et al.*[11] |
| **(CDRα) S51M** | N.P.[b] | N.P.[b] | -0.37 | Haidar *et al.*[11] |
| **(CDRα) S29A** | N.P.[b] | N.P.[b] | 0.33 | Haidar *et al.*[11] |
| **(CDRα) S100Y** | N.P.[b] | N.P.[b] | 2.34 | Haidar *et al.*[11] |
| **(CDRα) S100T** | N.P.[b] | N.P.[b] | -0.49 | Haidar *et al.*[11] |
| **(CDRα) S100N** | N.P.[b] | N.P.[b] | 1.92 | Haidar *et al.*[11] |
| **(CDRα) S100A** | N.P.[b] | N.P.[b] | -0.01 | Haidar *et al.*[11] |
| **(CDRα) R27F** | N.P.[b] | N.P.[b] | -0.22 | Haidar *et al.*[11] |
| **(CDRα) Q30N** | N.P.[b] | N.P.[b] | 0.73 | Haidar *et al.*[11] |
| **(CDRα) Q30E** | N.P.[b] | N.P.[b] | 0.43 | Haidar *et al.*[11] |
| **(CDRα) K68H** | N.P.[b] | N.P.[b] | 1.09 | Haidar *et al.*[11] |
| **(CDRα) G28V** | N.P.[b] | N.P.[b] | 0.56 | Haidar *et al.*[11] |
| **(CDRα) G28T** | N.P.[b] | N.P.[b] | -0.76 | Haidar *et al.*[11] |
| **(CDRα) G28R** | N.P.[b] | N.P.[b] | 2.43 | Haidar *et al.*[11] |
| **(CDRα) G28M** | N.P.[b] | N.P.[b] | -0.51 | Haidar *et al.*[11] |
| **(CDRα) G28L** | N.P.[b] | N.P.[b] | -0.46 | Haidar *et al.*[11] |
| **(CDRα) G28I** | N.P.[b] | N.P.[b] | -0.27 | Haidar *et al.*[11] |
| **(CDRα) G28A** | N.P.[b] | N.P.[b] | 0.58 | Haidar *et al.*[11] |
| **(CDRα) D26W** | N.P.[b] | N.P.[b] | -1.08 | Haidar *et al.*[11] |
| **(CDRα) D26V** | N.P.[b] | N.P.[b] | 1.45 | Haidar *et al.*[11] |
| **(CDRα) D26M** | N.P.[b] | N.P.[b] | 0.19 | Haidar *et al.*[11] |
| **(CDRβ) R102Q** | N.P.[b] | N.P.[b] | 0.45 | Haidar *et al.*[11] |
| **(CDRβ) I54R** | N.P.[b] | N.P.[b] | 1.28 | Haidar *et al.*[11] |
| **(CDRβ) G101A** | N.P.[b] | N.P.[b] | -1.09 | Haidar *et al.*[11] |
| **Data set 2**[a] | | | | |
| **WT** | 3200 | -7.50 | 0 | Cole *et al.*[12] |
| **A6c134M** | 1900 | -7.81 | -0.31 | Cole *et al.*[12] |
| **A6c134S** | 1800 | -7.84 | -0.34 | Cole *et al.*[12] |
| **A6c134AE** | 9.4 | -10.96 | -3.46 | Cole *et al.*[12] |
| **A6c134E** | 4.4 | -11.41 | -3.91 | Cole *et al.*[12] |
| **A6c134R** | 8 | -11.06 | -3.55 | Cole *et al.*[12] |
| **A6c134** | 4 | -11.47 | -3.96 | Cole *et al.*[12] |

[a] For both studies, $\Delta\Delta G$ values were determined using the WT affinity determined in the same study.

[b] N.P. means not provided in the original publication (only $\Delta\Delta G$ values were provided).

**Table S4.** CDR Loop sequences for the 1G4 set of TCRs. WT positions subjected to mutagenesis are underlined. Mutations are colored in red and in bold. The unique extra proline mutation in the CDR3α of NY-6 alongside the different CDR3α loop sequence of NY-33 and NY-33A are highlighted in yellow.

| TCR I.D. | CDR2α 50-54 | CDR3α 94-103 | CDR2β 50-53 | TCRβF3 56-70 | CDR3β 94-97 |
|---|---|---|---|---|---|
| **WT** | IQSSQ | PTSGGSYIPT | GAGI | QGEVPNGYNVSRSTT | YVGN |
| **122** | I**SPW**Q | P**LLD**G**T**YIPT | **AIQT** | QGEVPNGYNVSRST**I** | YVG**D** |
| **121** | I**SPW**Q | P**LLD**G**T**YIPT | **AIQT** | QGEVPNGYNVSRST**I** | YVGN |
| **120** | I**TPW**Q | P**LLD**G**T**YIPT | **AIQT** | QGEVPNGYNVSRST**I** | YVG**D** |
| **119** | I**TPW**Q | P**LLD**G**T**YIPT | **AIQT** | QGEVPNGYNVSRST**I** | YVGN |
| **113** | I**TPW**Q | P**LLD**G**T**YIPT | **AIQT** | **R**GEVPNGYNVSRST**I** | Y**L**GN |
| **112** | I**SPW**Q | P**LLD**G**T**YIPT | **AIQT** | **R**GEVPNGYNVSRST**I** | Y**L**GN |
| **107** | I**SPW**Q | P**FT**GG**G**YIPT | **AIQT** | QGEVPNGYNVSRSTT | YVGN |
| **6** | IQSSQ | **H**TS**N**G**YFP**PT | GAG**T** | **R**GEVPNGYNVSRST**I** | Y**L**GN |
| **33** | IQSSQ | P**YQS**G**H**Y**M**PT | GAG**T** | **R**GEVPNGYNVSRST**I** | Y**L**GN |
| **33A** | IQSSQ | P**YQS**G**H**Y**M**PT | GAG**T** | QGEVPNGYNVSRST**I** | Y**L**GN |
| **c49c50** | I**PFW**Q | PTSGGSYIPT | **SVGM** | QGEVPNGYNVSRSTT | YVGN |
| **c49wt** | I**PFW**Q | PTSGGSYIPT | GAGI | QGEVPNGYNVSRSTT | YVGN |
| **c52c50** | I**SPW**Q | PTSGGSYIPT | **SVGM** | QGEVPNGYNVSRSTT | YVGN |
| **c53c50** | I**TPW**Q | PTSGGSYIPT | **SVGM** | QGEVPNGYNVSRSTT | YVGN |
| **c55c50** | I**MGH**Q | PTSGGSYIPT | **SVGM** | QGEVPNGYNVSRSTT | YVGN |
| **c56c50** | I**MGT**Q | PTSGGSYIPT | **SVGM** | QGEVPNGYNVSRSTT | YVGN |
| **wtc50** | IQSSQ | PTSGGSYIPT | **SVGM** | QGEVPNGYNVSRSTT | YVGN |
| **wtc51** | IQSSQ | PTSGGSYIPT | **AIQT** | QGEVPNGYNVSRSTT | YVGN |

**Table S5.** Calculated mean absolute deviations (MADs) in kcal mol$^{-1}$ for the linear fits of $\Delta\Delta G_{calc}$ *vs.* $\Delta\Delta G_{exp}$ for the 1G4 and A6 test sets. MADs are determined in the presence and absence of the outliers described in the main text (see also **Figure S3**, **Figure S4**). MADs determined here are for MMPB/GBSA calculations performed without any explicit water molecules present and without any entropy corrections.

| | 1G4 | 1G4 (No Outliers) | A6 | A6 (No Outliers) |
|---|---|---|---|---|
| **MMGBSA** | 4.28 | 2.83 | 1.49 | 1.20 |
| **MMPBSA Di 1** | 7.15 | 6.26 | 3.17 | 2.57 |
| **MMPBSA Di 2** | 4.02 | 3.12 | 2.01 | 1.55 |
| **MMPBSA Di 4** | 2.55 | 1.66 | 1.74 | 1.42 |
| **MMPBSA Di 6** | 2.29 | 1.45 | 1.71 | 1.47 |
| **MMPBSA Di 8** | 2.39 | 1.62 | 1.78 | 1.52 |

**Table S6.** Calculated mean absolute deviations (MADs) in kcal mol$^{-1}$ for the linear fits of $\Delta\Delta G_{calc}$ *vs.* $\Delta\Delta G_{exp}$ for the 1G4 and A6 test sets. MADs are determined in the presence and absence of the outliers described in the main text (see also **Figure S3**, **Figure S4**). MADs determined here are for MMPB/GBSA calculations performed with a varying number of explicit water molecules present, but without any entropy corrections added.

| | 1G4 | 1G4 No Outliers | A6 | A6 No Outliers |
|---|---|---|---|---|
| **MMGBSA No Waters** | 4.28 | 2.83 | 1.49 | 1.20 |
| **MMGBSA 10 Waters** | 4.75 | 3.30 | 1.94 | 1.31 |
| **MMGBSA 20 Waters** | 5.73 | 4.37 | 2.86 | 2.02 |
| **MMGBSA 30 Waters** | 5.42 | 3.95 | 2.94 | 1.81 |
| **MMGBSA 50 Waters** | 5.48 | 3.65 | 3.32 | 2.19 |
| **MMPBSA Di 6 No Waters** | 2.29 | 1.45 | 1.71 | 1.47 |
| **MMPBSA Di 6 10 Waters** | 2.76 | 2.04 | 1.84 | 1.65 |
| **MMPBSA Di 6 20 Waters** | 3.21 | 2.61 | 2.12 | 1.85 |
| **MMPBSA Di 6 30 Waters** | 3.34 | 2.79 | 2.34 | 2.03 |
| **MMPBSA Di 6 50 Waters** | 3.99 | 3.40 | 2.62 | 2.30 |

**Table S7.** Calculated mean absolute deviations (MADs) in kcal mol$^{-1}$ for the linear fits of $\Delta\Delta G_{calc}$ *vs.* $\Delta\Delta G_{exp}$ for the 1G4 test set in the presence of different and solute entropy correction methods. MADs are determined in the presence and absence of the outliers described in the text (see also **Figure S3**). MADs determined here are for MMPBSA calculations performed with an internal dielectric constant of 6 and in the presence of either 0 or 50 explicit water molecules present.

| | **1G4** | **1G4 (No Outliers)** |
|---|---|---|
| **No Entropy** | | |
| **MMPBSA Di 6 No Waters** | 2.39 | 1.62 |
| **MMPBSA Di 6 50 Waters** | 3.99 | 3.40 |
| **Trunc NMA** | | |
| **MMPBSA Di 6 No Waters** | 2.22 | 1.97 |
| **MMPBSA Di 6 50 Waters** | 2.63 | 2.34 |
| **Int-Entropy** | | |
| **MMPBSA Di 6 No Waters** | 3.72 | 2.41 |
| **MMPBSA Di 6 50 Waters** | 4.73 | 3.47 |

(9) Li, Y.; Moysey, R.; Molloy, P. E.; Vuidepot, A. L.; Mahon, T.; Baston, E.; Dunn, S.; Liddy, N.; Jacob, J.; Jakobsen, B. K.; Boulter, J. M. Directed Evolution of Human T-

Cell Receptors with Picomolar Affinities by Phage Display. *Nat. Biotechnol.* **2005**, *23*, 349–354.

(10) Dunn, S. M.; Rizkallah, P. J.; Baston, E.; Mahon, T.; Cameron, B.; Moysey, R.; Gao, F.; Sami, M.; Boulter, J.; Li, Y.; Jakobsen, B. K. Directed Evolution of Human T Cell Receptor CDR2 Residues by Phage Display Dramatically Enhances Affinity for Cognate Peptide-MHC without Increasing Apparent Cross-Reactivity. *Protein Sci.* **2006**, *15*, 710–721.

(11) Haidar, J. N.; Pierce, B.; Yu, Y.; Tong, W.; Li, M.; Weng, Z. Structure-Based Design of a T-Cell Receptor Leads to Nearly 100-Fold Improvement in Binding Affinity for PepMHC. *Proteins Struct. Funct. Bioinforma.* **2009**, *74*, 948–960.

(12) Cole, D. K.; Sami, M.; Scott, D. R.; Rizkallah, P. J.; Borbulevych, O. Y.; Todorov, P. T.; Moysey, R. K.; Jakobsen, B. K.; Boulter, J. M.; Baker, B. M.; Li, Y.; Yi Li. Increased Peptide Contacts Govern High Affinity Binding of a Modified TCR Whilst Maintaining a Native PMHC Docking Mode. *Front. Immunol.* **2013**, *4*, 1–8.
